# Medial temporal lobe functional network architecture supports sleep-related emotional memory processing in older adults

**DOI:** 10.1101/2023.10.27.564260

**Authors:** Miranda G. Chappel-Farley, Jenna N. Adams, Richard F. Betzel, John C. Janecek, Negin S. Sattari, Destiny E. Berisha, Novelle J. Meza, Hamid Niknazar, Soyun Kim, Abhishek Dave, Ivy Y. Chen, Kitty K. Lui, Ariel B. Neikrug, Ruth M. Benca, Michael A. Yassa, Bryce A. Mander

## Abstract

Memory consolidation occurs via reactivation of a hippocampal index during non-rapid eye movement slow-wave sleep (NREM SWS) which binds attributes of an experience existing within cortical modules. For memories containing emotional content, hippocampal-amygdala dynamics facilitate consolidation over a sleep bout. This study tested if modularity and centrality—graph theoretical measures that index the level of segregation/integration in a system and the relative import of its nodes—map onto central tenets of memory consolidation theory and sleep-related processing. Findings indicate that greater network integration is tied to overnight emotional memory retention via NREM SWS expression. Greater hippocampal and amygdala influence over network organization supports emotional memory retention, and hippocampal or amygdala control over information flow are differentially associated with distinct stages of memory processing. These centrality measures are also tied to the local expression and coupling of key sleep oscillations tied to sleep-dependent memory consolidation. These findings suggest that measures of intrinsic network connectivity may predict the capacity of brain functional networks to acquire, consolidate, and retrieve emotional memories.

## INTRODUCTION

Our memories are important. They make us who we are, painting a picture of our past and guiding our future actions. The mechanisms that govern this process—determining which memories endure while others fade—have been a major topic of investigation for decades. Since the days of Ebbinghaus,^1^ who first sought to measure memory behaviorally, the field has learned much about the neurobiological substrates of memory processing.

A wealth of research has established that memories containing emotional content persist much longer than non-emotional experiences,^2^ and that medial temporal lobe (MTL) structures, such as the hippocampus and amygdala, play important roles in the acquisition, consolidation, and retrieval of emotional memories.^3–8^ The hippocampus is known to encode unique memory representations via pattern separation—the neural process of differentiating highly similar experiences as distinct events—which allows for the storage of similar experiences without overwriting previous ones. Amygdala activity is known to modulate hippocampal representations of emotional content,^9^ and the dynamic interplay between these MTL subregions is thought to be largely responsible for the preferential consolidation of emotional memories.^9–12^ While the precise mechanism underlying this process has long been theorized, accumulating evidence points to an important role for sleep in this process.

David Marr was the first to propose that memory consolidation occurred during offline periods of waking rest or sleep.^13^ During sleep, connectivity patterns in the brain are thought to reorganize to optimize information storage and integrate newly acquired knowledge into existing networks. While Donald Hebb first suggested that the modification of mutually connected synapses within cell assemblies supports learning,^14^ synaptic connectivity is extremely sparse; simply modifying existing synaptic weights would be insufficient for memory acquisition. To account for this, the cortex could organize into a hierarchical modular structure so that sparse connections between groups of highly interconnected regions (i.e., modules) would be sufficient to support memory encoding and storage.^15^ The hippocampus would serve as an index^16^ capable of coordinating activity patterns in modules representing different attributes of an experience.^15^ Through repeated reactivations of the hippocampal index, it is theorized that reinstatement of relevant activity patterns would strengthen intermodular connections (i.e., increase network integration) to facilitate memory consolidation. Notably, this reactivation and rearrangement of connections is proposed to occur during non-rapid eye movement Stage 3 (N3) sleep [i.e., slow-wave sleep (SWS)].^13,15,17,18^ Many theoretical frameworks have provided different accounts for the precise process by which initially labile memory traces are transformed into long-term representations,^19–21^ and while these theories may diverge in the degree to which they theorize hippocampal involvement in recall, all generally agree that hippocampal activity and its influence over network organization are necessary for memory acquisition and consolidation. Altogether, these theories suggest that a more modular network structure and hippocampal influence over this topological organization support memory consolidation, particularly for that which occurs during sleep.

Network neuroscience techniques, and specifically graph theoretic measures, have only recently started to be applied to study memory function.^22^ These tools provide a unique and innovative approach to test the theoretical premise of memory consolidation. Consolidation is theorized to involve reactivation of the hippocampal index code, which reinstates relevant patterns of activity stored in cortical modules which, over time, become reciprocally connected.^15^ Findings from the field of network neuroscience suggest that the human brain exhibits a modular organization in which subnetworks (i.e., modules) serve specific cognitive functions.^22–26^ Graph theoretical approaches conceptualize these modules as groups of highly interconnected nodes (anatomically defined regions) and edges (resting-state functional connectivity between regions).^23^ Network modularity (Q) is a global index of the modular organization of brain networks.^23,24,26^ Indeed, this intrinsic measure of network organization has been related to various facets of cognitive function,^27^ including episodic memory encoding^28^ and retrieval.^29^ However, whether Q supports memory consolidation, particularly over a sleep bout, has yet to be examined. Graph theory can also be used to investigate the topological roles of individual nodes. Centrality measures, such as eigenvector (EC) and betweenness centrality (BC), capture the relative import of individual nodes and their role in information flow.^30,31^ Taken together, these two metrics—when applied to the hippocampal formation—are akin to a hippocampal index; these measures identify the level of influence the hippocampus exerts over functional network topology (i.e., lower-level modules) and the degree to which it mediates information propagation, as proposed by hippocampal indexing theory and the hippocampus serving to reinstate patterns of activity associated with learning. However, it is unknown whether these measures of hippocampal centrality are associated with sleep-related memory processing.

Using these approaches, the current study aimed to address these knowledge gaps by combining high-resolution resting-state functional magnetic resonance imaging (fMRI) data, with overnight in-lab sleep polysomnography (PSG), and pre- and post-sleep memory testing taxing emotionally modulated hippocampal pattern separation. Two primary hypotheses were tested in a sample of cognitively healthy older adults: (1) that greater Q would be associated with increased NREM SWS expression and better overnight emotional memory retention, and (2) that greater hippocampal EC would be associated with better overnight emotional memory retention. Given the known involvement of the amygdala in emotional memory consolidation,^12,32^ this study also aimed to dissociate the contributions of the hippocampus and amygdala to each phase of emotional memory processing using BC as a secondary set of analyses. In a subset of participants, this study also explored whether these graph metrics were associated with the frontal expression of slow oscillations and frontal slow oscillation-sleep spindle coupling, which are tied to overnight memory retention and thought to reflect hippocampal-neocortical dialogue during sleep. Finally, random forest classification was used to establish the predictive power of these graph theoretical metrics for emotional memory encoding, consolidation, and retrieval.

## METHODS

### Study Participant Details

Forty cognitively healthy older adults (Mini-Mental State Exam score between 25-30) between the ages of 60-85 years from the BEACoN cohort were recruited for the overnight sleep study. All participants were free of major neurological and psychiatric disorders and were not taking any medications known to affect sleep or cognition, including non-SSRI antidepressants, neuroleptics, chronic anxiolytics, or sedative hypnotics. Participants had not traveled across time zones >3h time shift within at least 3 weeks prior to participation and did not have any diagnosis of sleep disorders other than sleep-disordered breathing (SDB) prior to participating; those with prior diagnoses were not being actively treated with continuous positive airway pressure. Participants did not consume any caffeine-containing food or beverages any time after 9AM on the day of their sleep study. Participants were fluent in English, had normal or corrected-to-normal vision and hearing, and were free of MRI contraindications. All participants provided written informed consent according to procedures approved by the Institutional Review Board of the University of California, Irvine (UCI). Of the initial 40 overnight sleep study participants, three subjects were excluded due to fMRI FOV misalignment and one subject declined MRI scanning, leaving data from 36 participants for analyses.

### Emotional Mnemonic Discrimination Task

Participants performed the emotional version of the mnemonic discrimination task (eMDT) prior to and following overnight sleep (see **Figure 1**).^9^ The eMDT is an established framework to measure hippocampal-dependent emotional memory, specifically pattern separation processes reliant on interactions between the DG/CA3 subregions of the hippocampus and the amygdala.^9,33^

**Figure 1.**
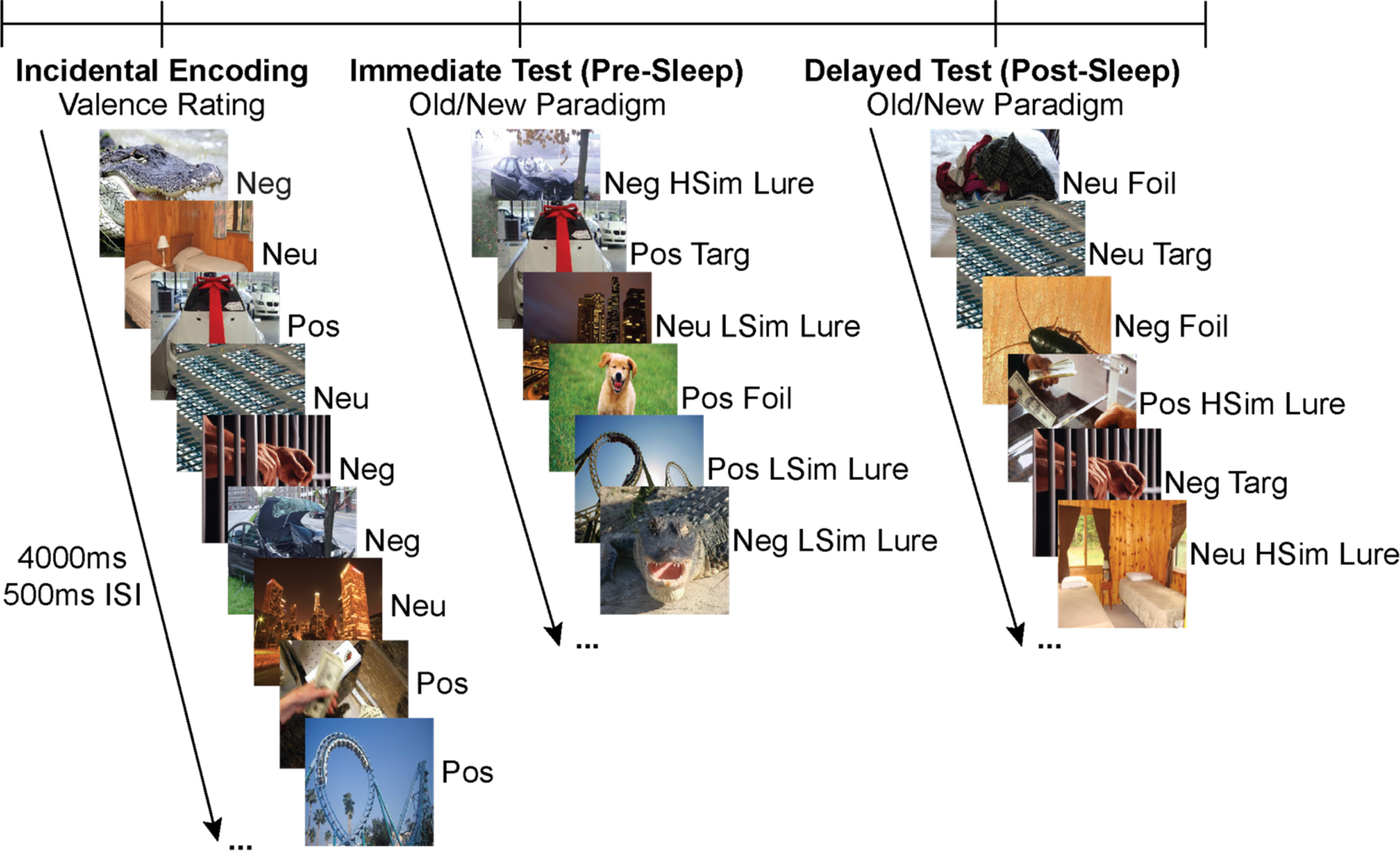
Emotional Mnemonic Discrimination Task. The eMDT consists of three phases: incidental encoding and a surprise immediate test phase prior to sleep, and a delayed test phase following overnight sleep. During the incidental encoding phase, participants are shown a series of emotionally salient images and asked to rate them as positive, negative, or neutral via button press. Following encoding, participants perform an old/new recognition paradigm and are shown targets, foils, and high/low similarity lures modulated by valence. Following overnight sleep, participants are again shown targets, foils, and lures and are asked to indicate via button press whether it is an old or new image. Images from the incidental encoding phase are randomly split into the immediate and delayed test phases. ISI—Inter Stimulus Interval; Neg—Negative; Neu—Neutral; Pos—Positive; LSim—Low Similarity; HSim—High Similarity; Targ—Target

A Windows laptop equipped with Python 2.7 and PsychoPy v1.90.3 was used to present the task stimuli and record participant responses. Each trial consisted of an image presentation screen alternating with a fixation cross. Each image was presented on a black background for 4 seconds, which was followed by a fixation display consisting of a white fixation cross on a black background for 500ms. Responses were recorded using 3 individual colored orby button switches (P.I. Engineering, Williamston, MI).

Participants underwent an incidental encoding phase, in which they were shown 180 emotionally salient images presented in a random order. They rated each image on valence (i.e., positive, negative, or neutral) via a button press while the image was present. Immediately following the encoding phase, participants underwent a surprise memory test (immediate test phase), wherein they were presented with another 120 images presented in random order. These images were split evenly amongst exact repeats (30 targets), new images (30 foils), and images similar to targets (60 total lures). Lure images were either highly similar to targets (30 high similarity; HSim) or somewhat similar to targets (30 low similarity; LSim). Stimuli were binned *a priori* for emotional valence and similarity.^9^ While each image was present, participants indicated via button press whether they had seen that exact image before (‘old’) or whether it was an entirely new image (‘new’). Following overnight sleep, participants completed another surprise memory test (delayed test), wherein they were presented with 120 images split evenly amongst targets, foils, and lures as outlined above. No targets or corresponding lures presented in the immediate test phase were shown during the delayed test phase. Each valence and similarity type were distributed evenly between targets, foils, and lures.

Performance was calculated using the bias-corrected Lure Discrimination Index (LDI), defined as p(‘new’|lures) – p(‘new’|Targets). LDI was calculated collapsing across similarity bins for each valence condition at both immediate and delayed testing timepoints. Overnight memory retention was measured via overnight change in LDI (LDI_delayed test_-LDI_immediate test_). Analyses were initially *a priori* restricted to LDI calculated for negative stimuli for two main reasons: (1) the positive and negative stimuli in this task are not matched on arousal,^33^ and (2) prior work has demonstrated that negative elements of an experience are preferentially consolidated during sleep,^34–36^ whereas positive components of an experience do not necessarily show a sleep benefit.^34^ However, to show specificity of the relationships between network measures and the consolidation of emotional stimuli, secondary exploratory analyses were conducted using LDI metrics for neutral images. Unless noted otherwise, analyses were conducted using LDI calculations collapsing across interference conditions (HSim + LSim lures). However, given established literature that older adults generally require greater dissimilarity to effectively pattern separate, negative LSim LDI (low interference condition) was explored in relation to Q when a relationship with negative LDI was not present.

### Polysomnography and Sleep Architecture

Electroencephalography (EEG) with polysomnography (PSG), including electrooculogram (EOG) electromyogram (EMG), electrocardiogram (ECG), and respiratory monitoring were recorded during overnight sleep using an integrated PSG-EEG system (Natus software v9.1.1) and ANT Neuro Waveguard gel cap (either 128 or 256 channels). EEG was collected at 1024 Hz and downsampled to 256 Hz. Sleep scoring was visually staged in 30-s epochs and events were marked by Registered Polysomnographic Technologists according to the American Academy of Sleep Medicine guidelines^37^ and verified by a board-certified sleep physician (RMB). Main measures derived from the overnight sleep study included the percentage of time spent in each sleep stage, total sleep time, sleep efficiency (total sleep time/ total time in bed), and the Apnea-Hypopnea Index (AHI), which is a commonly used metric capturing the severity of SDB. While the majority of participants reported no prior diagnosis of SDB during study enrollment, the prevalence of SDB increases with age^38^ and frequently goes undiagnosed.^39^ Therefore, AHI was included as a covariate in all statistical models. However, some reports suggest that AHI may be an insufficient measure to capture the complexity of SDB and its impacts on neurobiology and behavior^40^ given that it only captures the average frequency of apneas and hypopneas per hour. Therefore, time spent below 90% oxygen saturation was derived as a secondary measure of SDB-related hypoxic burden.

### EEG Preprocessing and Slow Wave Spectral Analysis

Spectral analyses were performed on a subset of participants (N=18) with data collected from the 128-channel EEG caps. Analyses focused on this subset of participants in order to analyze the highest quality data, as data collection methods were switched to the 128-channel cap following data quality issues with the 256-channel cap. Additionally, the two nets were in different sensor space, making it difficult to combine accurately localized data across participants. While data from all electrodes was preprocessed, a subset of 19 channels with a similar spatial distribution to the standard 10-20 system were selected to maximize signal quality from electrodes that were minimally dropped during the cleaning process (AFF1, AFF2, FC1, F3, FZ, F4, F6, FFC5H, C3, FCC1H, C6, FTT8H, CPP6H, P5, P3, POZ, T6, O1, P4). Given *a priori* hypotheses focusing on frontal slow oscillation expression, outcome measures were averaged across frontal derivations corresponding to electrodes AFF1, AFF2, FC2, F3, FZ, F4, and F6.

EEG preprocessing was performed using semi-automated artifact rejection^41–44^ with custom MATLAB (MathWorks, Inc, R2019b) scripts focusing on NREM sleep stages (N2 and N3 [N2N3]) with the EEGLAB toolbox (https://sccn.ucsd.edu/eeglab/index.php).^45^ First, data were imported into EEGLAB and down sampled to 256Hz. EEG data were then notch filtered (59.5-60.5Hz) and bandpass filtered in the 0.3Hz–35Hz frequency range, both using a 5500-point Hamming windowed sinc Finite Impulse Response (FIR) filter. Channels were average-referenced and manually inspected for noise; electrodes were dropped from analyses only if noise was present for a large proportion of the sleep period. Artifact rejection was performed in three steps.^41–44^ First, instances of arousals, apneas, and respiratory event-related arousals were automatically rejected while concatenating NREM stages N2 and N3. Second, epochs with high-powered artifact bursts were removed using an automated 99.8 percentile threshold based on broadband power spectral density. Third, epochs were visually inspected, and segments of noisy data or artifact-heavy electrodes were removed. Lastly, for electrodes dropped during preprocessing, spherical spline interpolation was used to reconstruct EEG signal from nearby derivations. Power spectral density estimates for slow-wave activity (SWA: 0.5-4 Hz), and slow oscillation (SO: 0.5-1 Hz) and delta (1-4 Hz) frequencies at each electrode were calculated with Welch’s method using a 6-second Hamming window. Absolute and relative power were calculated for each frequency of interest at each channel location and then averaged across the aforementioned frontal derivations to calculate absolute or relative frontal SWA, SO, and delta power.

### Slow Oscillation-Sleep Spindle Phase Amplitude Coupling

Slow oscillation (SO) sigma phase amplitude coupling (PAC) was calculated using a two-step procedure. First, SOs were detected from clean, artifact-free concatenated N2N3 sleep from the same 19 channels used in spectral analyses by first measuring zero-crossings of filtered EEG signal in the 0.5-1 Hz frequency range. Filtered EEG data with negative to positive zero crossings separated by 0.5 to 1.5 second were detected. The detected zero crossings were labeled as SOs if the following criteria were met: 1) the minimum amplitude from the SO trough to peak (i.e., peak to peak amplitude, 60 μV), 2) the minimum wave depth of the negative half of the SO (i.e., volt threshold, 30 μV), 3) the time between the first and second zero crossing (i.e.., SO duration, 300ms – 1.5s), and 4) the total duration of the event (i.e., distance between first and third zero crossing; SO period, 10s). While prior work in young adults has used a peak-to-peak threshold and volt threshold of 80 μV, ^46,47^ using these thresholds in the current study yielded no detected SOs, likely due to known age-related reductions in SO amplitude.^48–50^ Therefore, the current study used a peak-to-peak amplitude of 60 μV and volt threshold of 30 μV^50^, as a recent study found SOs of amplitudes as low as nearly 60μV in age groups similar to that of the current study cohort.^48^ Next, the SOs and fast sigma were Butterworth filtered (SO: 4^th^ order high and low Butterworth filter; fast sigma 8^th^ and 9^th^ order high and low Butterworth filter). The Hilbert transform was then applied to the filtered signals to compute the composite complex-valued signal yielding the amplitude of fast sigma and phase of the SO. The absolute value of the mean of this complex vector, termed the Modulation Index^51^—but is most similar to mean vector length (MVL) and will hereafter be referred to as such—corresponds to the coupling strength between slow oscillation phase and fast sigma amplitude. Non-zero values of MVL indicate coupling between SO phase and fast sigma amplitude. However, this raw value does not account for potential overestimates due to non-white noise in the signals. Therefore, MVL values were normalized by comparing our MVL values to those generated from surrogate data incorporating an arbitrary time lag between SO phase and fast sigma amplitude. A distribution of surrogate MVL values were computed across a range of different lags, with the normalized MVL computed as:

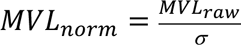

The normalized MVL was calculated for each of the 19 channels, and the average normalized MVL of the aforementioned frontal derivations was calculated derivations and used in analyses. Two subjects were excluded from these exploratory analyses due to the absence of sleep spindles or SOs (final N=16).

### MRI Acquisition

All neuroimaging data were collected at the Campus Center for Neuroimaging (CCNI) at the UCI, using a 3-Tesla Siemens Magnetom Prisma scanner equipped with a 32-channel head coil. The average time difference between the MRI scan and overnight sleep study visit was 1.3 ± 1.0 years. Structural MRI data were acquired using a T1-weighted MPRAGE sequence with the following parameters: orientation=sagittal, TR=2300ms, TE= 2.38ms, FA=8, voxel resolution=0.8mm isotropic, FOV=256mm. Participants had rsfMRI data which was obtained using a T2*-gradient echo planar imaging (EPI) sequence sensitive to blood-oxygenation level dependent (BOLD) contrast with 84 volumes acquired in 39 interleaved slices, with 1.8mm isotropic voxels: TR=2500ms, TE= 26ms, FA=70, FOV= 200mm. Of this sample, 2 participants had slightly longer scans with 240 volumes, and 3 participants had scans with 360 volumes, TR=1600ms, FOV = 212mm. For the purposes of analyses, these differences did not systematically affect signal-to-noise ratio.

### Resting-State Functional MRI Preprocessing

Neuroimaging data were batch preprocessed according to scan parameters using CONN toolbox^52^ version 21.a implemented in MATLAB 2019b and Statistical Parametric Mapping (SPM12, Wellcome Trust Centre for Neuroimaging, London, United Kingdom). Structural images were centered, segmented into gray matter, white matter, and cerebrospinal fluid components, and normalized to the Montreal Neurological Institute (MNI) template (1mm^3^). The functional images were centered and aligned to the AC-PC line, volumes were realigned and unwarped, slice-time corrected to their specific TR, and normalized to the MNI template (2mm^3^). Following recommendations for network and ROI-based analysis,^53^ functional data were not smoothed to maintain integrity of distinct BOLD signal time series across smaller, adjacent ROIs. Outlier volumes were detected using Artifact Detection Tools (ART) implemented in CONN using a threshold of motion >0.5mm/TR and a global intensity z-score of 3. Denoising consisted of the six realignment parameters and their first-order derivatives, spike regressors generated from outlier detection,^54^ and anatomical CompCor (first five components from white matter and CSF). Preprocessed resting-state scans for each participant were linearly detrended and bandpass filtered (0.008–0.09 Hz) was applied after nuisance regression (RegBP). A threshold of >20% volumes detected as outliers was used to flag subjects with high residual motion for exclusion. However, we found that subjects had minimal motion (average framewise displacement for valid scans, μ=0.06±0.03) and all subjects survived this threshold.

To determine ROIs for inclusion in analyses, an explicit mask of coverage across the partial FOV of the functional images was created. Using FSL (v1.0.13), this explicit mask was applied to the Brainnetome Atlas^55^ to determine which ROIs were present across all subjects. This atlas was specifically chosen because it is a functional parcellation based on known anatomical boundaries which provides smaller ROIs consistent with our high resolution MTL focused scan, which allowed us to maximize the number of ROIs within the somewhat limited FOV. Any region with less than 50% coverage across all subjects was excluded, resulting in 151 ROIs with sufficient coverage within the FOV for all participants (see **Figure 2, Tables S1-S2**). Mean BOLD timeseries were extracted after denoising from each of the 151 Brainnetome ROIs for ROI-to-ROI first-level analyses in CONN. This generated a 151 x 151 functional connectivity matrix for each subject, reflecting the Fisher’s r-to-z transformed correlation coefficients for each pair of ROIs, used for graph analyses.

**Figure 2.**
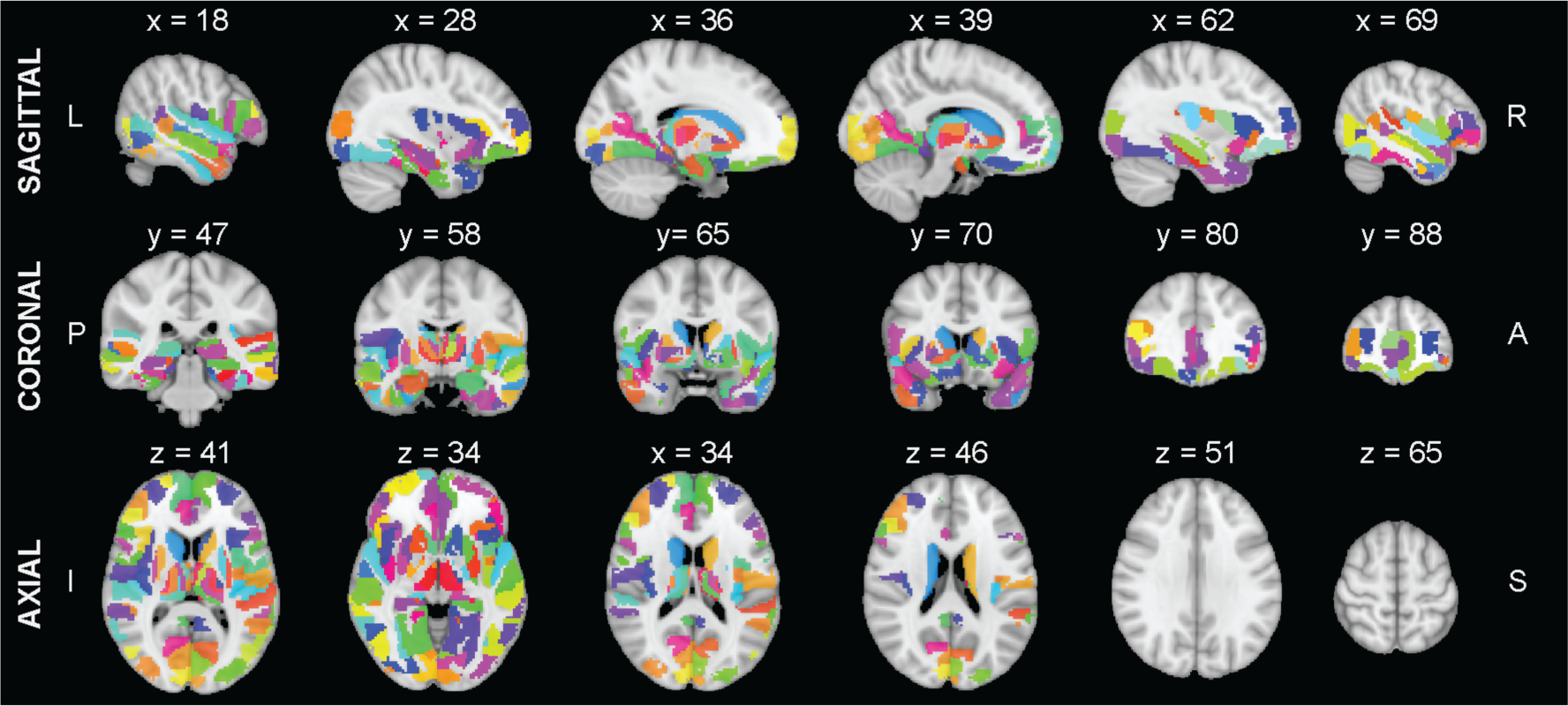
Regions of Interest (ROIs) included in the analysis mask for all subjects.

### Graph Theory Analyses

The Brain Connectivity Toolbox (BCT), implemented in MATLAB 2021a, was used for all graph analyses.^56^ ROI-to-ROI connectivity matrices were optimized (ensured matrix symmetricity, diagonals set to zero, normalized) to generate weighted, signed adjacency matrices for each subject for graph analyses. A signed and weighted approach was used rather than the positive-only, binary, cost-modified approach to network construction as weak or negative correlations may still be neurobiologically relevant, particularly when investigating individual differences.^26,57^ Three graph theoretical measures were primarily used for analyses: network modularity (Q), eigenvector centrality (EC), and betweenness centrality (BC). Participation coefficient (PC) was derived as a secondary measure for analysis.

#### Network Modularity (Q)

Q captures the degree to which a network can be divided into communities, or modules. Higher Q indicates a more modular network structure. Q was computed using the “community_louvain” function in BCT by running the Louvain Modularity Maximization algorithm using a uniform null model^58^ 150 times to generate consensus partitions across 7 different resolution parameters (γ) ranging from 0 (assuming zero correlation between nodes) to 2 standard deviations away from the mean positive connection weight in 0.05 steps (see **Figure S2**).^59^ The uniform null model was selected to account for the transivity of the input matrix and for ease of interpretation, as nodes are only included in the same community if their connectivity exceeds the given γ-value.^59^ This approach allowed us to test the consistency of results when resolving larger (smaller γ-values) and smaller (larger γ-values) community structures to ensure our findings were not specific to a given resolution parameter. The final Q value used in analyses corresponds to Q derived from the consensus partition at γ equal to the median positive connection weight (0.10).

#### Eigenvector Centrality (EC)

EC is a measure of the influence a node exerts over a network that accounts for both the quantity and quality of connections by considering the degree of the node of interest, and the degree of its neighbors.^30^ Greater EC suggests that a given node may be highly influential, based on how influential its neighbors are as well. EC was derived using the “eigenvector_centrality_und” function in BCT on the optimized adjacency matrix. Negative weights were remapped prior to computation.^30^ Average EC was calculated across all hippocampus nodes (left/right, rostral/caudal) to generate an average hippocampal EC value, as well as across all amygdala nodes (left/right, medial/lateral) to generate a value for average amygdala EC. Average EC of the parahippocampal cortex (BNA ROI A35/36: left/right, rostral/caudal), occipital region (BNA ROI vmPOS: left/right), and medial frontal gyrus (A10l: left/right) were also derived and used as both proximal and distal control regions to demonstrate specificity of effects.

#### Betweenness Centrality (BC)

BC is an index of a node’s ability to mediate information propagation based on shortest paths.^31^ High BC indicates that a node is ‘central’ to information flow through a network, as many shortest paths go through that node to reach the rest of the network. Lower BC may suggest that a given node is less functionally connected, and thereby less important for directing information flow. To calculate BC, adjacency matrices were optimized by first dropping negatives and then remapping edge weights to lengths using the BCT function “weight_conversion.” BC values were calculated from the positive, length matrix using the function “betweenness_wei” and normalized to [0,1]. Average BC was calculated across all hippocampus nodes (left/right, rostral/caudal) to generate an average hippocampal BC value, as well as across all amygdala nodes (left/right, medial/lateral) to generate a value for average amygdala BC. Average BC of the parahippocampal cortex, occipital, and medial frontal regions were also derived for control analyses.

#### Participation Coefficient (PC)

PC measures the distribution of a node’s connections across network communities. Greater PC (values closer to 1) suggests that a node’s connections are evenly distributed across modules, whereas lower values (closer to zero) suggest that a node’s connections are more localized to a singular community.^26^ To calculate PC, the final partitions generated from Modularity Maximization were used to calculate the positive PC at each resolution parameter using the function “participation_coef_sign” in BCT. The PC calculated at γ equal to the median positive connection weight (0.10) was used for control analyses.

### Statistical Analyses

All analyses were conducted in IBM SPSS v27, RStudio v2022.12.0+353, or in Python v3.8.5. Assumptions for multiple linear regression were checked and several variables were transformed to meet assumptions of normality: AHI was log transformed and a constant was added (+1) to avoid log of zero, PC measures were reflected (2-x) and log transformed, and amygdala, parahippocampal, and medial frontal gyrus BC were cube root-transformed. Prior to analyses, Kendall’s rank correlation was calculated between mean motion calculated from all valid MRI scans and each graph metric to ensure network measures were not driven by motion. No statistically significant associations were found between mean motion and each primary network measure used in analyses (see **Table S3**). It is also important to note that no statistically significant associations between the graph measures and either AHI or SDB-related hypoxic burden were observed (**Table S4**). All analyses included self-reported biological sex as a binary covariate, and age and log-transformed AHI as continuous covariates unless noted otherwise. Models were run initially with these three covariates, but as a follow-up, analyses were performed a second time including the time difference between the overnight sleep study and MRI scan as an additional covariate.

A multiple regression framework adjusting for the covariates mentioned above was used to test each primary hypothesis with Q, EC, and BC as outlined in the results. For all analyses with Q, sensitivity analyses were performed to ensure that findings were consistent across resolution parameters (see **Table S5, Figure S3**). Given recent research demonstrating that SDB impacts SWS expression,^60^ associations between AHI (Pearson’s r) and time spent below 90% oxygen saturation (Kendall’s τ_b_) were additionally explored (**Table S4)**. Hayes SPSS Process Macro (v4.2)^61^ was used to test for an indirect effect of Q on overnight change in negative LDI through NREM SWS expression (model 4), adjusting for the covariates mentioned above. This generated 95% percentile-based bootstrapped confidence intervals (CIs) based on 5000 samples. When upper and lower bootstrapped CIs did not include 0, the indirect effect was considered statistically significant at the α=0.05 level. Sensitivity analyses were performed to ensure that this mediation effect was consistent across resolution parameters (**Figure S3**). As much research suggests that REM sleep is important for emotional memory consolidation^62^, a control multiple regression model using REM sleep percentage as the IV and overnight change in negative LDI as the DV while adjusting for the same covariates was also tested. To show specificity of results to stimuli of negative valence, control regression models using LDI for neutral stimuli, adjusting for the same covariates, were also constructed. Relationships between graph measures and sleep oscillatory activity were exploratory in nature, and therefore were tested initially with Pearson’s correlation. In the case of findings with delayed retrieval, multiple - regression models were initially re-run adjusting for pre-sleep memory performance as a proxy for overnight consolidation. Follow-up analyses were then performed adjusting for age and sex, unless otherwise noted, due to the limited sample size for these analyses.

Given that both hippocampal and amygdala EC were associated with overnight emotional memory retention, regression commonality analysis (CA) was conducted to parse apart individual and common effects of hippocampal and amygdala EC in predicting overnight change in negative LDI. CA decomposes regression variance into unique and common effects of each independent variable^63^, thereby enabling us to determine the degree to which hippocampal and amygdala EC contribute distinct (accounted for by each single predictor) or shared information (accounted for by both predictors) to overnight change in negative LDI. The effects of each covariate were first regressed out of the IVs (hippocampal and amygdala EC) and DV (overnight negative memory retention) and the proportional share effects were calculated with CA in RStudio using the ‘yhat’ package.^64^

To identify the most important graph theoretic metrics and determine their utility in predicting each stage of memory processing, Random Forest (RF) classification using leave-one-out cross validation (LOOCV) was performed. RF is a supervised machine learning algorithm that builds many decision trees from a random sample of training data for classification based on majority voting. RF using LOOCV was specifically chosen due to its ability to handle multidimensional data and smaller sample sizes.^65^ First, class variables were generated from the continuous performance measures using median split, such that a 1 or 0 for negative LDI at immediate or delayed test phases would indicate better/worse discrimination performance, whereas a 1/0 for overnight change in negative LDI would indicate better retention/more forgetting. RF models using LOOCV implemented with Python Scikit-Learn library were then generated.^66^ We built 100 decision trees initially consisting of different combinations of graph theoretic metrics of interest. The input features were Q, hippocampal EC, amygdala EC, hippocampal BC, and amygdala BC. Feature importance was evaluated using the Gini Impurity Index.^67^ Using the top three features, the algorithm was run again and its performance was evaluated using AUC of the ROC curves, which was plotted from the predicted values of the test data using LOOCV. Bootstrapped CIs for the ROC were calculated by resampling 1000 times with replacement using a 1.0 sampling rate.

## RESULTS

### Demographics and Protocol

Thirty-six participants (μ_age_=72.9 ± 5.6 years, 63.8% female, **Table 1**) recruited from the Biomarker Exploration in Aging, Cognition, and Neurodegeneration (BEACoN) cohort at the University of California, Irvine, completed in-lab PSG to derive measures of sleep architecture and oscillatory activity. An emotional mnemonic discrimination task (eMDT) was completed during the overnight sleep study visit with the encoding and immediate test phases performed prior to PSG recording and delayed testing occurring the following morning (see **Figure 1**). Task performance was assessed using the Lure Discrimination Index (LDI) which is thought to be a behavioral proxy for hippocampal pattern separation.^9,68,69^ The measure of overnight consolidation was derived as the change score from the delayed (post-sleep) and immediate (pre-sleep) timepoints. See **Table 2** for a summary of behavioral data across valence categories; memory performance in this sample was consistent with prior work in a sample of cognitively intact older adults.^70^ Analyses were restricted *a priori* to stimuli of negative valence, as prior research suggests that negative aspects of an experience are primed for preferential consolidation during sleep.^34–36^ However, follow-up exploratory analyses were performed with images of neutral valence to demonstrate specificity to emotional stimuli. As part of the BEACoN study, these participants completed high-resolution structural and resting-state fMRI at a separate timepoint which was used for graph analyses and to construct measures of intrinsic network structure and the trait-like topological roles of specific MTL nodes.

**Table 1.** Participant Characteristics.

| **Demographics** | **N= 36** |
| --- | --- |
| Age, Mean (SD) | 72.9 (5.6) |
| Sex, No. Female (%) | 23 (63.9) |
| Years of Education, Mean (SD) | 16.6 (2.3) |
| MMSE, Mean (SD) | 28.6 (1.5) |
| BMI, Mean (SD) | 24.5 (5.8) |
| AHI, Mean (SD) | 12.0 (14.3) |
| Race, No. |  |
| African American or Black | 1 |
| Asian | 9 |
| White | 23 |
| More than one Race | 2 |
| Unknown/Not Reported | 1 |
| Ethnicity, No. |  |
| Hispanic | 1 |
| Non-Hispanic | 35 |
| Sleep Architecture |  |
| Total Sleep Time (min), Mean (SD) | 342.6 (65.8) |
| Sleep Efficiency, Mean (SD) | 77.0 (14.0) |
| N1 (min), Mean (SD) | 33.8 (18.5) |
| N2 (min), Mean (SD) | 190.3 (35.2) |
| N3 (min), Mean (SD) | 57.8 (34.4) |
| REM (min), Mean (SD) | 60.8 (29.3) |
| WASO (min), Mean (SD) | 96.3 (65.2) |

Abbreviations: MMSE—Mini Mental State Exam; BMI—Body Mass Index; AHI—Apnea-Hypopnea Index; NREM—Non-rapid eye movement sleep; REM—Rapid eye movement sleep; WASO—Wake after sleep onset

**Table 2.**
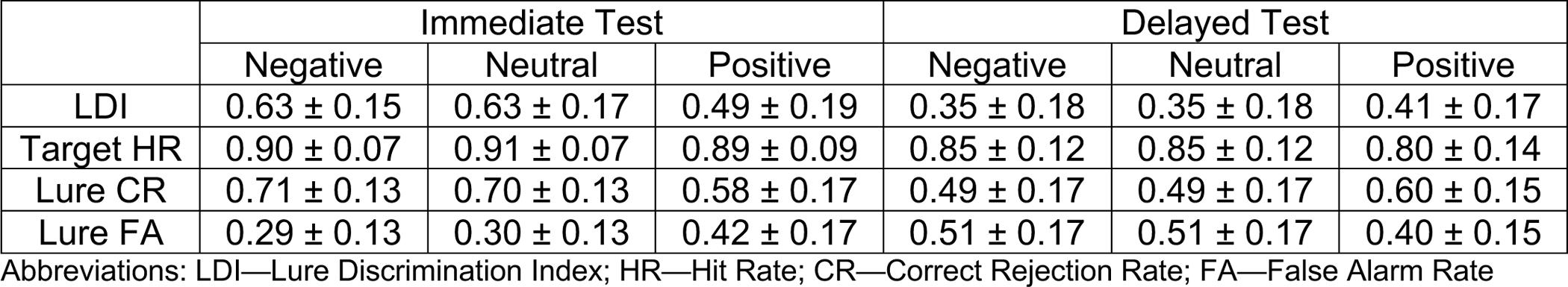
Behavioral data from the E-MDT.

### Characterizing resting-state network properties with graph theory

To examine overall functional network composition as well as the topological roles of the hippocampus and amygdala, three graph theoretic metrics were derived: (1) modularity (Q; **Figure 3a**), which measures the degree to which brain activity is segregated into modules that are sparsely connected to the rest of the network;^22,24,26^ (2) hippocampal/amygdala eigenvector centrality (EC; **Figure 5a**), which measures the influence of these nodes by accounting for both the quantity and quality of functional connections;^30^ and (3) hippocampal/amygdala betweenness centrality (BC; **Figure 6a**), which captures the extent to which these nodes may be important for information flow within a network based on shortest paths (i.e., strongest functional connections).^31^ To compute these metrics, the average BOLD timeseries for each subject was extracted from 151 Brainnetome Atlas^55^ regions of interest (ROIs) included within the restricted FOV of the MTL-focused high-resolution resting-state scan (**Figure 2; Table S1-S2**). ROI-to-ROI connectivity matrices were generated for each subject, symmetrized, and normalized to generate weighted adjacency matrices for computation of Q, EC, and BC.

**Figure 3.**
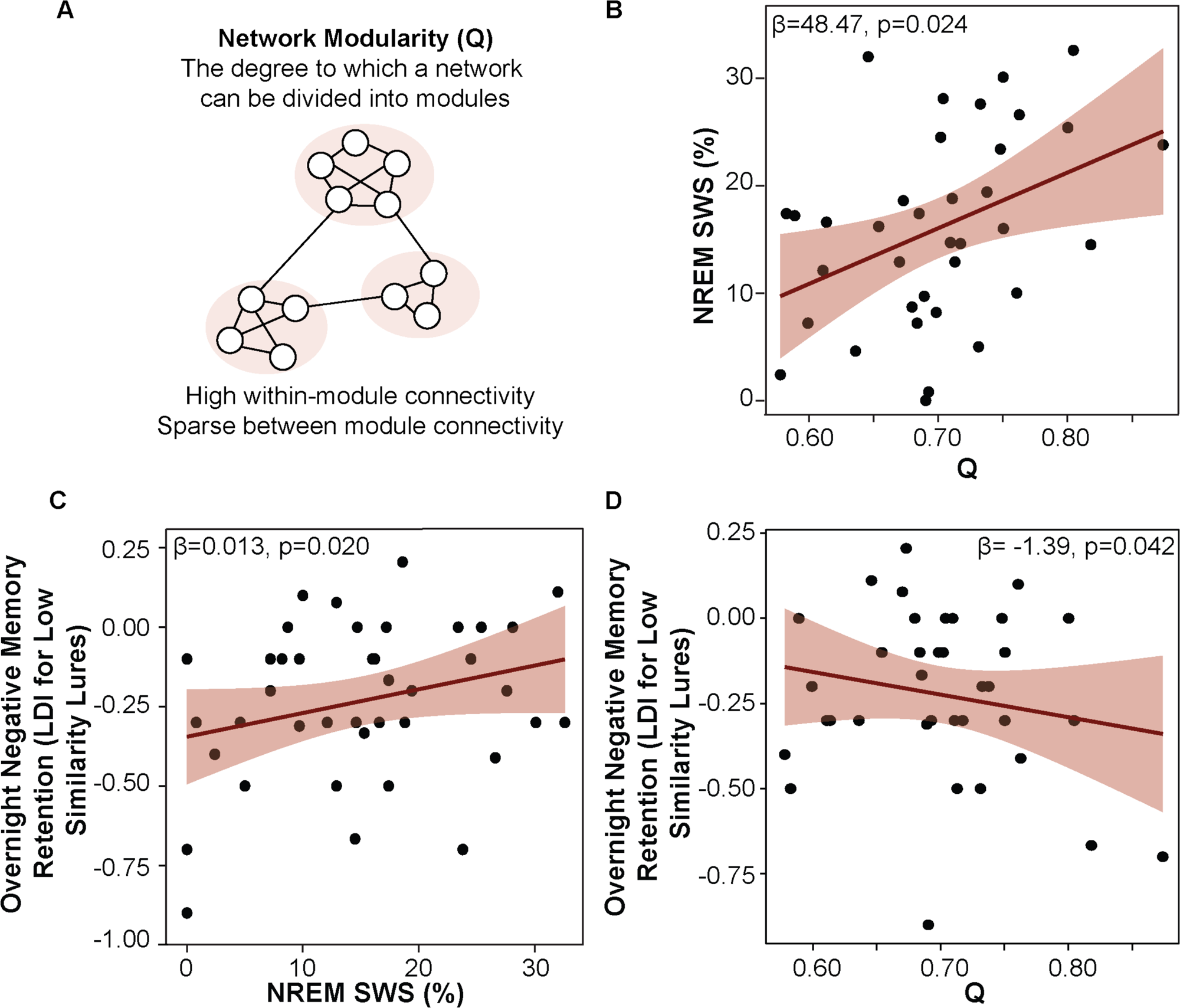
More modular brain networks express more NREM SWS sleep and exhibit worse overnight memory retention. (A) Schematic of network modularity. (B) Greater network modularity is associated with a greater percentage of sleep time spent in NREM SWS, while adjusting for age, sex, and AHI. (C) A greater percentage of sleep time spent in NREM SWS is positively associated with overnight emotional memory retention, adjusting for age, sex, AHI, and Q. (D) Greater network modularity is negatively associated with overnight emotional memory retention, adjusting for age, sex, AHI, and percentage of sleep time spent in NREM SWS.

### NREM SWS mediates the relationship between network modularity and emotional memory retention

Q is a network-wide measure that evaluates the degree to which a network can be divided into modules, with modules being groups of densely interconnected nodes that are sparsely connected to the rest of the network.^26^ Q was calculated across all network nodes based on an optimization procedure with 150 repetitions; the final Q used in analyses was generated from the consensus partition calculated at the resolution parameter (γ-value) corresponding to the median connection weight (see Methods). High Q values are generally indicative of a more segregated network structure, whereas lower values of Q may suggest a more integrated network topology.

First, we examined whether Q (from MRI collected at a separate timepoint) was associated with sleep architecture. NREM SWS was analyzed due to its theorized role in memory consolidation and reactivation. Multiple regression analysis with NREM SWS percentage as the dependent variable (DV), Q as the independent variable (IV), and age, self-reported biological sex, and AHI as covariates revealed that a more modular network structure was associated with a greater percentage of time spent in NREM SWS (β=48.476, SE=20.343, p=0.023; **Figure 3b**). Sensitivity analyses demonstrated consistency in this result across resolution parameters (γ-values), which are incorporated into the optimization algorithm to tune the number and size of detected modules (**Table S3**). Although recent work suggests sleep-disordered breathing (SDB) impacts SWS expression,^60^ no statistically significant associations between either AHI (p=0.773) or hypoxic burden (i.e., time below 90% oxygen saturation; p=0.142) and NREM SWS were observed in the current study. These data suggest that more modular brain networks, in general, express more NREM SWS, a period of sleep associated with a breakdown of local functional connectivity^71^ and enhanced network integration.^72,73^

Next, the relationship between NREM SWS and overnight emotional memory retention was tested independent of its relationship with Q. A multiple regression analysis using NREM SWS percentage as the DV, overnight change in negative LDI as the IV, and age, sex, AHI, and Q as covariates revealed no statistically significant relationship (p=0.292). However, it is well established that older adults require greater dissimilarity among inputs (i.e., lower interference) to effectively pattern separate.^68^ Therefore, we examined the relationship between NREM SWS and LDI which was calculated using only low similarity negative lures (negative LSim LDI). A multiple regression model using NREM SWS as the DV, overnight change in negative LSim LDI as the IV, adjusting for the same covariates, revealed a statistically significant association between NREM SWS and overnight retention of emotional memories (β=0.013, SE=0.005 p=0.020; **Figure 3c**). NREM SWS was not associated with overnight retention of neutral memories (overnight change in neutral LSim LDI, p=0.318). Together, these findings suggest that NREM SWS may support overnight emotional memory retention.

Then, the relationship between Q and overnight emotional memory retention was tested independent of its relationship with NREM SWS. A model using overnight change in negative LSim LDI as the DV, Q as the IV, and age, sex, AHI, and NREM SWS as covariates revealed a significant negative relationship between Q and overnight retention of negative stimuli (β=-1.394, SE=0.655, p=0.042; **Figure 3d**), suggesting that trait-like network integration (i.e., lower Q from MRI collected at a separate timepoint) supports sleep-related processing of emotional memories under low levels of interference (LSim condition). A control multiple regression model confirmed the specificity of this effect to negative stimuli (overnight change in neutral LSim LDI as the DV, p=0.893). A summary of behavioral data under low levels of interference (LSim condition) can be found in **Table S6**.

Finally, to formally test whether NREM SWS mediated the association between Q and overnight retention of emotional memories, ordinary least squares path analyses were conducted using overnight change in negative LSim LDI as the DV, Q as the IV, NREM SWS as the mediating variable, and age, sex, and AHI as covariates. A conceptual diagram of the mediation model is shown in **Figure 4**.

**Figure 4.**
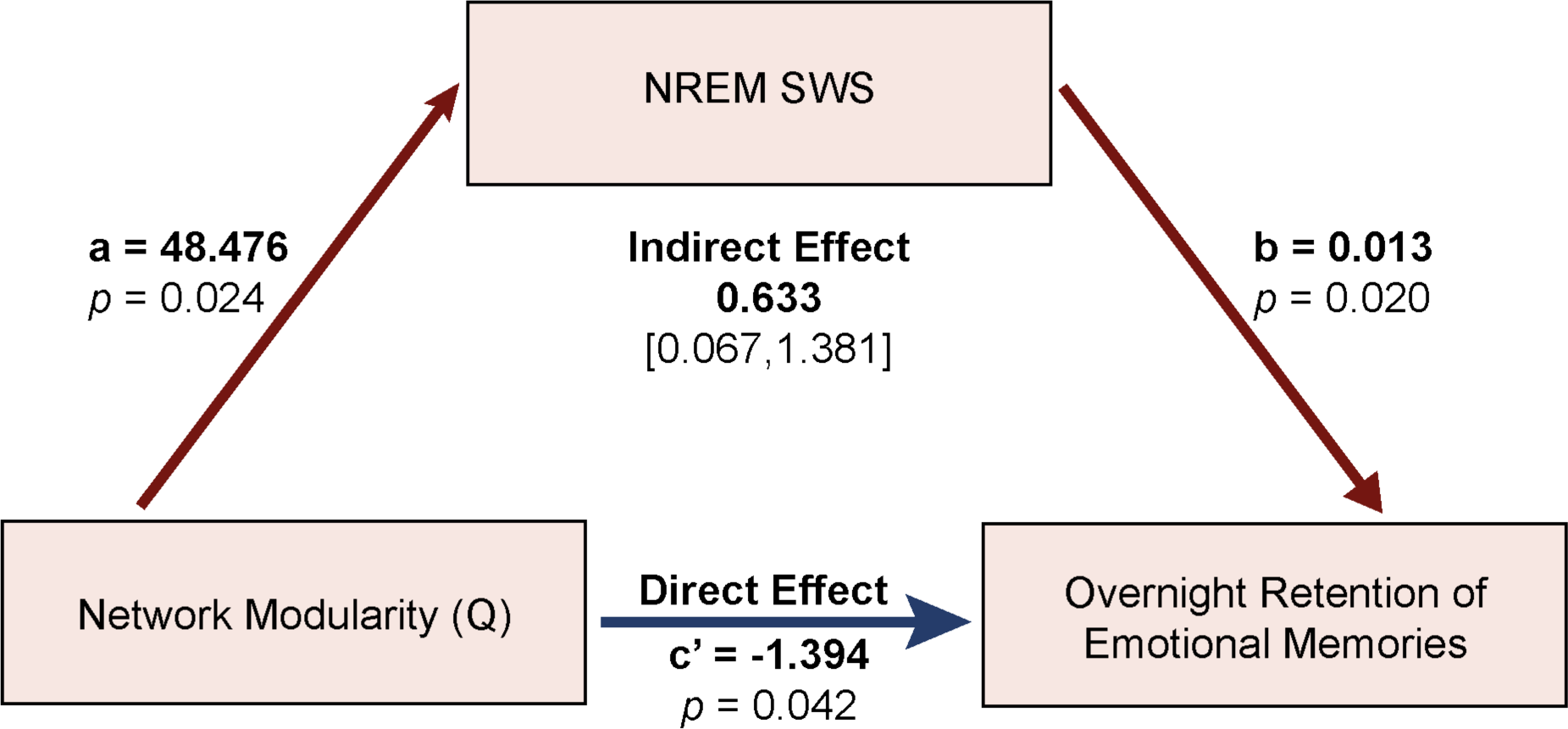
NREM SWS mediates the relationship between network modularity and emotional memory retention. NREM SWS percentage of sleep time is shown as a mediator of the relationship between network modularity and overnight emotional memory retention.

Greater Q was associated with more NREM SWS (a=48.476, standardized a= 0.373, SE=20.243, p=0.024), which, in turn, was associated with better overnight retention of negative stimuli (b=0.013, standardized b= 0.471, SE=0.005, p=0.020). Percentile-based bootstrapped 95% confidence intervals (CIs) for the indirect effect did not include zero (0.633, 95% CIs: 0.067, 1.381; standardized indirect effect: 0.176, 95% CIs: 0.015, 0.345). After adjusting for NREM SWS, the direct effect of Q on overnight negative memory retention remained significant, suggesting a partial inconsistent mediation by NREM SWS in this relationship (c’=-1.394, standardized c’ = −0.388, SE=0.655, p=0.042). Sensitivity analyses confirmed that this mediation effect held across a range of resolution parameters aside from the highest γ-value (see **Figure S3**). Importantly, when the IV and mediating variable were swapped, Q did not significantly mediate the relationship between NREM SWS and overnight emotional memory retention. A separate mediation model accounting for the time between the MRI scan and overnight sleep study in addition to the covariates also supported a significant indirect effect of Q on overnight emotional memory retention via NREM SWS (0.6506, 95% CIs: 0.043, 1.451). While much research implicates REM sleep in emotional memory processing,^62^ a multiple regression model using REM sleep percentage as the IV instead of NREM SWS as a control analysis did not reveal a statistically significant association between REM and overnight change in negative LSim LDI (p=0.759). These results may indicate that greater trait-like network integration either directly, or indirectly—perhaps during SWS—supports overnight emotional memory retention (see Discussion for further interpretation).

### Hippocampal-amygdala functional network hierarchy supports emotional memory retention

EC is a graph metric that derives node importance within a functional network by accounting for both the *quantity and quality* of connections, such that higher EC indicates a node is strongly connected to other strongly connected nodes.^30^ To determine whether resting-state hippocampal EC predicts the degree of overnight emotional memory retention, a multiple regression model was constructed using overnight change in negative LDI as the DV, hippocampal EC as the IV, and age, sex, and AHI as covariates. This analysis revealed a significant positive association between hippocampal EC and overnight emotional memory retention (β=36.845, SE=12.993, p=0.008; **Figure 5a**). This result remained statistically significant when including the time difference between the MRI scan and overnight sleep study as an additional covariate (p=0.017). Hippocampal EC was not associated with overnight change in neutral memories (p=0.544). This suggest that a more influential hippocampus, as measured by both the quantity and quality of its connections (EC), aids in overnight emotional memory preservation.

**Figure 5.**
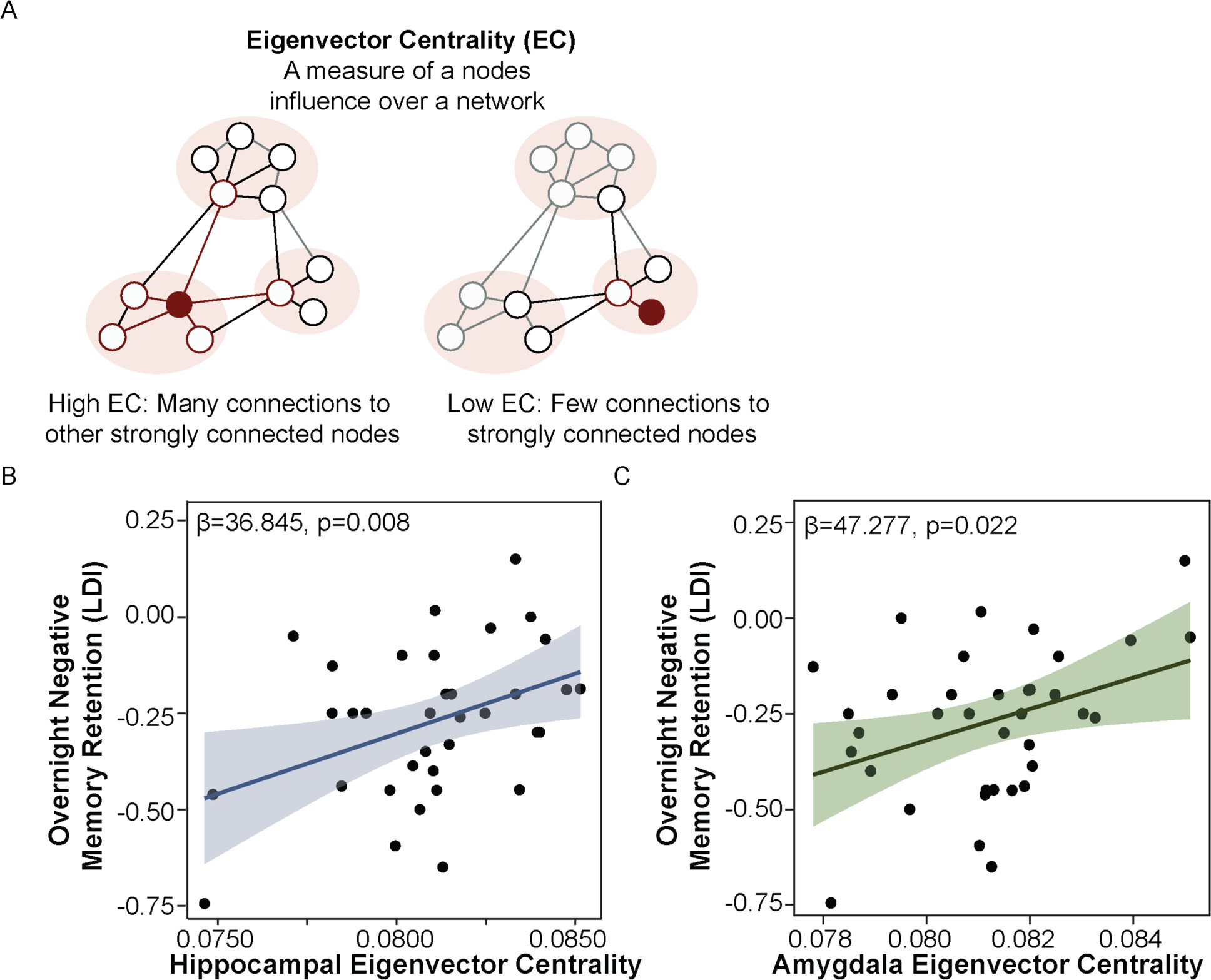
Hippocampal-amygdala functional network hierarchy supports emotional memory retention. (A) Schematic of eigenvector centrality. (B) Greater hippocampal eigenvector centrality is positively associated with overnight emotional memory retention, adjusting for age, sex, and AHI. (C) Greater amygdala eigenvector centrality is positively associated with overnight emotional memory retention, adjusting for age, sex, and AHI.

As a control analysis, we tested these relationships with participation coefficient (PC), which measures the distribution of a node’s connections across communities and is greatest when a node’s connections are uniformly distributed across communities.^26^ This metric differs from EC, which accounts for the *quality* of a node’s connections in addition to the quantity. Therefore, to determine whether it was the positioning of the hippocampus within the resting-state functional network hierarchy (i.e., the fact that the hippocampus is strongly functionally connected to other strongly functionally connected nodes or modules) or simply the distribution of its intermodular connections (i.e., quantity of connections across modules) that was important for overnight emotional memory retention, a separate multiple regression model was constructed using hippocampal PC as the IV instead of hippocampal EC, while adjusting for covariates. Importantly, PC was not significantly associated with overnight change in negative LDI (p=0.682), suggesting that it is the influence of the hippocampus over network organization that supports overnight emotional memory retention, rather than simply the distribution of its connections across network modules.

Given the known involvement of the amygdala in task performance^9,33^ and the role of hippocampal-amygdala dynamics in emotional memory consolidation, it was necessary to determine whether amygdala resting-state EC also contributes to overnight emotional memory retention. A multiple regression model predicting overnight change in negative LDI from amygdala EC, adjusting for the same covariates mentioned above, demonstrated a significant positive association between amygdala EC and overnight emotional memory retention (β=47.277, SE=19.553, p=0.022; **b**). This result remained statistically significant when also adjusting for the time difference between the MRI scan and sleep study (p=0.016). Amygdala EC was not related to overnight neutral memory retention (p=0.613).

To tease apart hippocampal and amygdala contributions to overnight emotional memory retention, a follow-up analyses was performed to examine the interaction effect between hippocampal and amygdala EC in predicting overnight change in negative LDI while adjusting for age, sex, and AHI. This analysis revealed no significant interaction between hippocampal and amygdala EC (p=0.793). To determine the unique and shared contributions of hippocampal and amygdala resting-state EC to overnight emotional memory retention, regression commonality analysis (CA) was performed. After first regressing out the effects of the covariates on both predictors (hippocampal and amygdala EC) and the outcome measure (overnight change in negative LDI), CA revealed that hippocampal EC accounts for roughly 40% of the variance in overnight emotional memory retention, whereas amygdala EC accounts for roughly 22% of the variance. CA also indicated that roughly 38% of the variance is common to both amygdala and hippocampus EC (see **Table 3**). These findings may suggest that intrinsic hippocampal-amygdala dynamics are at least as important as the unique contributions of each MTL node for the capacity to retain emotional information overnight, if not more so as in the case of the amygdala.

**Table 3.** Results from Commonality Analysis.

|  | **Coefficient** | **% Total Variance** |
| --- | --- | --- |
| Unique to Hippocampal EC | 0.1049 | 39.81 |
| Unique to Amygdala EC | 0.0576 | 21.87 |
| Common to Both | 0.1010 | 38.31 |

Abbreviations: EC—eigenvector centrality

To determine whether it was the topological influence of these nodes rather than their shared connectivity that is important for overnight emotional memory retention, a multiple regression model was constructed to test the association between hippocampal-amygdala resting-state functional connectivity and overnight emotional memory retention while adjusting for the covariates. This analysis revealed a trending association between greater hippocampal-amygdala functional connectivity and overnight change in negative LDI (β=0.399, SE=0.226, p=0.087). This may indicate that the influence of these MTL nodes over resting-state network architecture, and their common topological dynamics (revealed by CA), rather than their connectivity strength, is most important for overnight emotional memory retention.

As a final set of control analyses, multiple regression models testing whether EC of proximal MTL (parahippocampal) and distal cortical (ventromedial parietooccipital sulcus, medial frontal gyrus) regions were associated with overnight emotional memory retention were constructed. These areas were specifically selected to demonstrate specificity to hippocampal and amygdala MTL nodes and to ensure that these relationships were not a global brain-wide phenomenon. No significant associations between the EC of these regions and overnight change in negative LDI (all p’s > 0.200) were observed. Taken together, these results highlight the specificity of the hippocampus and amygdala in the resting-state functional network hierarchy in supporting overnight retention of emotional memories.

### Betweenness centrality identifies distinct roles for hippocampus and amygdala in emotional memory acquisition, consolidation, and delayed retrieval

While EC is a metric representing a nodes influence over a network, BC describes the relative importance of an individual node for information propagation *via* shortest paths (i.e., strongest connections). A node with high BC may be optimally functionally positioned to mediate information transfer through a network as it is strongly functionally connected to many, potentially disparate nodes (see **Figure 6a**).^74^ The CA results indicated that hippocampus and amygdala EC contribute unique, as well as shared, influence to facilitate the consolidation of emotional memories. Therefore, using BC, we sought to further delineate the roles of these two nodes in emotional memory acquisition, consolidation, and retrieval. Resting-state hippocampal BC was conceptualized as a measure of hippocampal memory processing akin to the hippocampal index which directs activity in lower-level modules during acquisition (i.e., strongly connected to many nodes and optimally functionally positioned to coordinate information propagation), and amygdala BC as amygdala modulation of emotional memory processing.^32^ Given this theoretical framework, we hypothesized that resting-state hippocampus BC would be associated with memory performance at immediate test whereas amygdala BC would be primarily associated with memory consolidation and retrieval.

**Figure 6.**
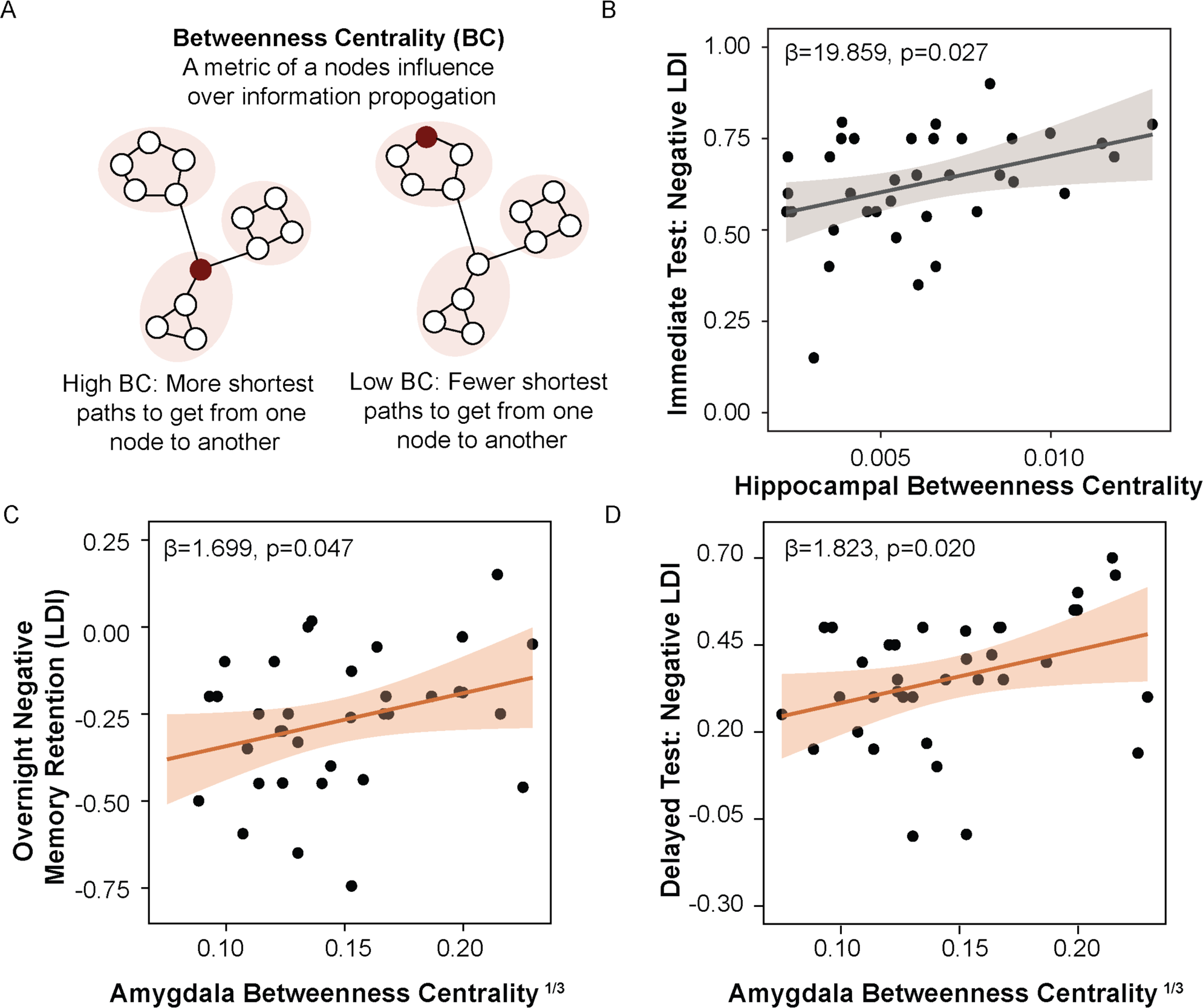
Betweenness centrality identifies distinct roles for hippocampus and amygdala in emotional memory acquisition, consolidation, and delayed retrieval. (A) Schematic of betweenness centrality. (B) Greater hippocampal betweenness centrality is positively associated with emotional memory performance at immediate test, adjusting for age, sex, and AHI. (C) Higher amygdala betweenness centrality (cube-root transformed) is positively associated with overnight emotional memory retention and (D) emotional memory performance at delayed test, adjusting for age, sex, and AHI.

To first test whether hippocampal BC predicted memory acquisition gauged by memory performance at immediate test, a multiple regression model using negative LDI at immediate test as the DV, hippocampal BC as the IV, and age, sex, and AHI as covariates was constructed. This analysis revealed that higher hippocampal BC was associated with better performance at immediate test for negative stimuli (β=19.859, SE=8.564, p=0.027; **Figure 6b**). Hippocampal BC was not related to immediate test performance of neutral stimuli (p=0.435). BC of the amygdala (p=0.855), parahippocampal (p=0.612), ventromedial parietooccipital sulcus (p=0.506), or medial frontal gyrus (p=0.551) ROIs also were not significantly associated with immediate test performance, showing specificity to the hippocampal formation.

To test whether amygdala BC was associated with overnight retention and next-day retrieval of emotional memories, multiple regression analysis was conducted using overnight change in negative LDI as the DV and amygdala BC as the IV, adjusting for the same covariates described above. This model revealed that higher amygdala BC was associated with better overnight retention of negative stimuli (β=1.699, SE=0.822, p=0.047; **Figure 6c**). A similar regression model, using negative LDI at delayed test as the DV, showed a positive association between amygdala BC and negative LDI (β=1.823, SE=0.745, p=0.020; **Figure 6d**). Amygdala BC was not associated with overnight neutral memory retention (p=0.616) or delayed test performance for neutral images (p=0.601). Hippocampal BC was not associated with overnight change in negative LDI (p=0.459), or negative LDI at delayed test (p=0.338). Control multiple regression analyses using parahippocampal, ventromedial parietooccipital sulcus, and medial frontal gyrus BC as the IVs and overnight change in negative LDI or negative LDI at delayed test as DVs further confirmed that these results were specific to the amygdala (all p’s ≥ 0.110)

Altogether, these results suggest that individual differences in hippocampal control over information flow (BC) during restful wake is important for emotional memory acquisition, as measured by performance after a short delay, whereas amygdala contributions to information propagation are particularly important for overnight consolidation and next-day retrieval of emotional information. Including the time difference between MRI scan and overnight sleep study as an additional covariate in analyses did not significantly impact these findings (all p’s ≤ 0.047).

### Random forest (RF) classification identifies most important resting-state graph features for sleep-related memory consolidation

Given results from multiple regression analyses which demonstrated that distinct resting-state graph theoretical metrics of overall network structure and nodal centrality were differentially tied to memory acquisition, consolidation, and delayed recall, RF classification, a type of supervised machine learning algorithm, was then used with LOOCV to determine the optimal combination of these measures for predicting each phase of memory processing. Receiver operating characteristic (ROC) curves in **Figure 7** demonstrate model performance of RF classification for overnight emotional memory retention (**Figure 7a**), delayed recall (**Figure 7b**), and immediate recall (**c**). The Gini Impurity Index was used to identify the most important resting-state graph theoretical metrics from all metrics for each stage of memory processing. Then, the top three features were selected to construct a final RF model using bootstrapping with 1000 simulations to calculate 95% CIs for area under the curve (AUC) for each ROC. The best prediction model for overnight emotional memory retention combined graph theoretical measures of amygdala BC, Q, and hippocampal EC (AUC = 0.77 ± 0.01, 95% CIs: 0.774 to 0.775). The best prediction model for delayed recall included graph measures of hippocampal EC, amygdala BC, and Q, with an AUC = 0.73 ± 0.02 (95% CIs: 0.731 to 0.734). The best prediction model for immediate test combined measures of Q, hippocampal BC, and hippocampal EC (AUC = 0.52 ± 0.03, 95% CIs: 0.522 to 0.525); however, this model had low predictive value nearly at chance levels. These results demonstrate that hippocampal and amygdala centrality measures, alongside resting-state functional network structure (Q), may have more moderate utility in predicting emotional memory consolidation than encoding.

**Figure 7.**
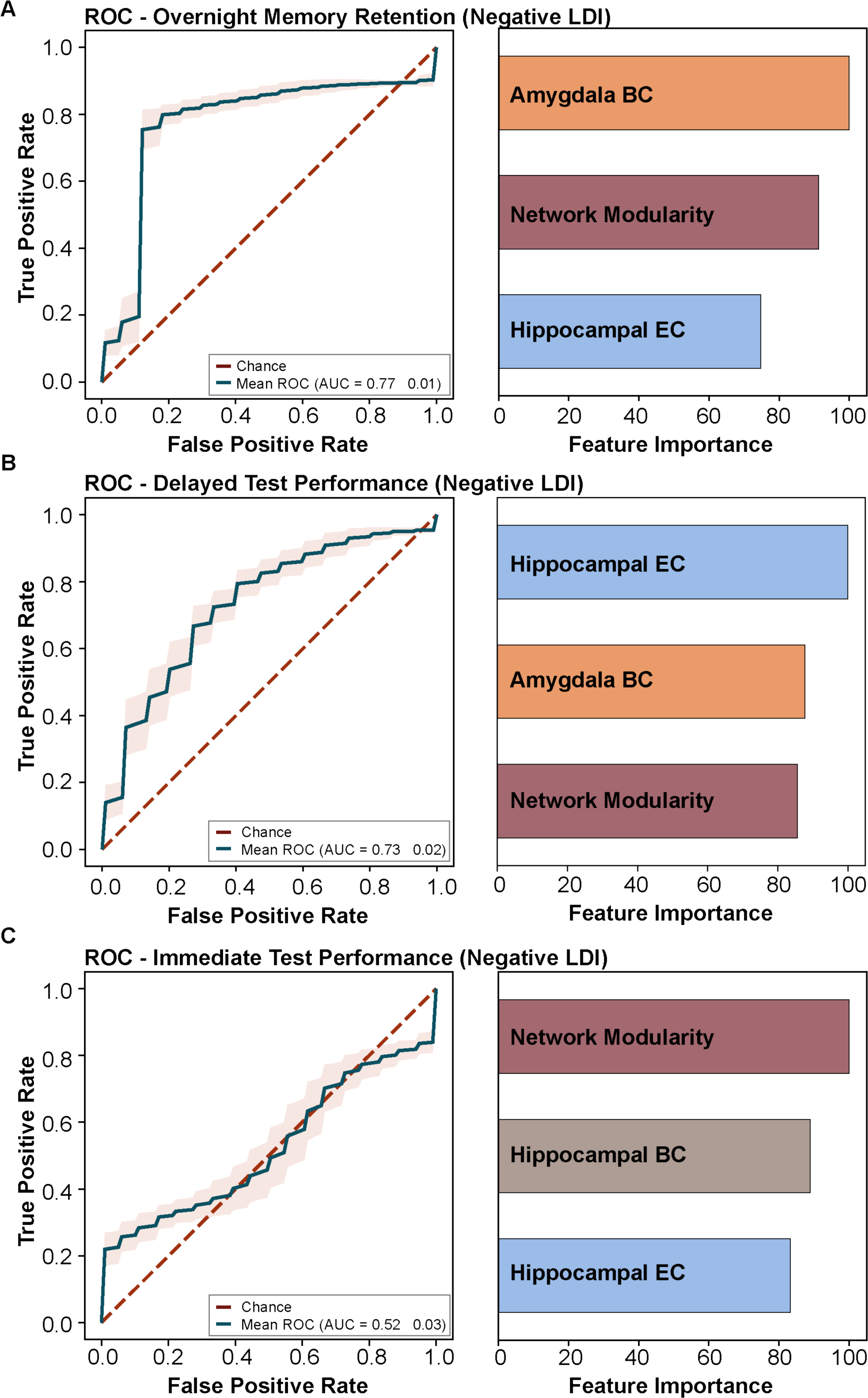
Random forest classification identifies most important graph features for sleep-related memory consolidation. (A) ROC and Gini Impurity Index output for top 3 predictors of overnight negative memory retention (LDI). (B) ROC and Gini Impurity Index output for delayed test performance for negative LDI. (C) ROC and Gini Impurity Index output for top 3 predictors of immediate test performance for negative LDI. ROC—Receiver Operating Characteristic Curve; AUC—Area Under the Curve; EC—Eigenvector Centrality; BC—Betweenness Centrality; LDI—Lure Discrimination Index

### Frontal slow oscillation expression is linked to emotional memory retention and measures of hippocampal centrality

Slow oscillations (SOs) are high amplitude (≥75µV), low frequency (0.5-1 Hz) waveforms primarily expressed during SWS that reflect neuronal firing synchrony, the spectral power of which is maximal over frontal EEG derivations.^75^ SOs are implicated in memory consolidation,^76^ and may facilitate long-range communication across brain regions during an otherwise dampened state of connectivity.^46^ Given findings with Q and NREM SWS which suggests that network integration supports overnight emotional memory consolidation, frontal SO power was explored as a more precise measure of underlying neural activity tied to memory processing.

While no statistically significant relationship was observed between frontal relative SO power and overnight emotional memory retention (p=0.293, N=18), frontal relative SO power was significantly associated with delayed retrieval (r=0.492, p=0.038, N=18; **Figure S4A**), which remained a trend when adjusting for pre-sleep memory performance (p=0.071), suggesting that SO power supports overnight emotional memory retention. However, this relationship did not survive adjustment for age and sex (p=0.237).

Despite statistically significant relationships between Q and SWS, Q was not significantly associated with frontal relative SO power (p=0.174, N=18). However, a correlation between SWS and frontal relative SO power approached statistical significance in the expected direction (r=0.458, p=0.056, N=18), perhaps suggesting that Q may still be related to greater frontal SO expression given that SWS is a period of sleep characterized by SOs.

Next, relationships between frontal SO expression and measures of hippocampal centrality were explored. Pearson’s correlation revealed a statistically significant association between hippocampal EC and frontal relative SO power (r=0.475, p=0.047, N=18; **Figure S4B**), which remained a trending predictor after adjusting for age and sex (β=4.180, SE=2.131, p=0.070). Hippocampal BC showed a trending correlation with frontal relative SO power (r=0.426, p=0.086, N=18; **Figure S4C**), which reached statistical significance when age and sex were added into the multiple regression model (β=3.77, SE=1.528, p=0.027). Interestingly, hippocampal EC was a strong predictor of emotional memory retrieval (r=0.552, p<0.001, N=18; **Figure S4D**). This relationship remained statistically significant when adjusting for pre-sleep performance as well as age and sex (β=42.537, SE=10.882 p<0.001). These exploratory analyses may suggest that brain networks with more influential hippocampi exhibit better overnight emotional memory retention both directly, and indirectly via greater relative SO expression.

### Measures of hippocampal and amygdala centrality are tied to frontal slow oscillation-sleep spindle coupling

Phase-amplitude coupling (PAC) between SOs and fast sleep spindles (SPs; 13-16 Hz) is thought to reflect synaptic changes that facilitate integration and reorganization of memory traces into existing networks via hippocampal-neocortical communication.^77–79^ SO-SP coupling was measured via mean vector length (MVL), which measures how tightly the phase providing (i.e., SO) and amplitude providing (i.e., fast SPs) signals are coupled, with higher values indicating strong coupling between SOs and fast SPs.^80^ We specifically focused on measures of MTL centrality, given hippocampal implication in memory transformation and amygdala modulation of consolidation processes. Hippocampal EC was significantly correlated with frontal SO-SP coupling (r=0.649, p=0.006, N=16**; Figure S5A**), which remained statistically significant after adjusting for age and sex (β=531.066, SE=206.867, p=0.025). Pearson’s correlation also revealed a statistically significant association between amygdala BC and frontal SO-SP coupling (r=0.545, p=0.029, N=16; **Figure S5B**). However, this relationship was no longer statistically significant after adjusting for age and sex (p=0.100). Frontal SO-SP PAC was not significantly associated with overnight emotional memory retention (p=0.477) or delayed retrieval (p=0.314). These data suggest that brain resting-state networks that exhibit more intrinsically influential hippocampi and greater amygdala modulation over information flow also exhibit greater frontal SO-SP coupling, a neural mechanism crucial for memory consolidation during sleep.

## DISCUSSION

Results from the current study suggest that intrinsic network properties (occurring during the resting-state) may statistically predict the capacity for brain functional networks to acquire, consolidate, and retrieve emotional memories, including over a night of sleep. These findings provide greater insight into how brain network functional systems operate in the context of memory processing, suggesting that more integrated subsystems (Q), and the topological influence of key MTL nodes over functional network architecture (EC) and activity (BC) are important for the encoding and sleep-related stabilization of emotional memories in older adults. A main finding is that greater network modularity was associated with NREM SWS expression, which, in turn, was associated with better overnight retention of emotional memories. These effects were specific to NREM SWS and were consistent across a range of resolution parameters. We further found that greater hippocampal and amygdala influence over network activity, measured by greater EC, also supported overnight consolidation of emotional memories. These effects were specific to the hippocampus and amygdala, and not proximal or distal control regions, and were not explained by their shared functional connectivity. Commonality analyses revealed the distinct and common contributions of each of these regions, demonstrating that both contribute unique and shared information. Hippocampal and amygdala BC were used as an index of their control over information propagation and these analyses identified roles for each of these MTL nodes in memory acquisition versus consolidation and retrieval. RF classification revealed that distinct combinations of these graph theoretical metrics exhibit potential predictive utility for memory consolidation and retrieval. Lastly, the relationships between graph measures and sleep oscillatory activity implicated in memory consolidation were explored. These analyses revealed that greater hippocampal influence over functional network organization is tied to greater frontal SO expression and SO-SP coupling. Collectively, these results provide support for two major tenets of memory consolidation theory: (1) that greater network integration, potentially during SWS, allows for the rearrangement of connections across network modules to support memory consolidation, and that (2) greater hippocampal influence over network structure and activity flow (akin to a hippocampal index) aids in the coordination of network activity and neural sleep oscillations to facilitate memory consolidation. The results from the current study also provide additional support for the importance of hippocampal-amygdala dynamics in sleep-related processing of emotional memories.

Greater resting-state network modularity was positively associated with the amount of time spent in NREM SWS, which was positively associated with overnight memory retention. However, an inconsistent mediation effect was observed. Inconsistent mediation occurs when the direct and indirect effect have opposite signs.^81^ In the present analyses, the direct effect linking network modularity to overnight emotional memory retention was negative, whereas the indirect effect revealed a positive effect of modularity on overnight memory retention through NREM SWS expression. While this result may initially seem counterintuitive, it is possible that both effects are reflecting the same neurobiological process. The direct effect suggests that reduced network modularity (i.e., greater network integration) is associated with better overnight emotional memory retention. The indirect effect indicates greater network modularity is associated with a greater percentage of time spent in NREM SWS—a period of sleep associated with a breakdown of brain network dynamics,^72^ cortical connectivity,^71^ and increased functional network integration^73^—and better overnight emotional memory retention. Therefore, these two effects, despite opposite signs, may be indicating that enhanced network integration either directly, or indirectly during NREM SWS expression, supports overnight memory retention. In fact, during NREM SWS, SOs have been shown to facilitate long-range communication and provide a window for integration which aids in the consolidation of memory.^46^ In further support of this notion, the current study also found a significant relationship between frontal relative SO power and overnight emotional memory retention. Altogether, these findings complement existing literature and theories. Prior work has demonstrated that increased network modularity supports episodic encoding,^28^ while a breakdown of modular structure during retrieval facilitates accurate recall.^29^ These works suggest that encoding relies upon a segregated, more modular brain state whereas successful recall relies upon a more integrated functional network structure. The present findings expand upon this literature and suggest that increased functional network integration (i.e., reduced modularity) supports overnight retention of emotional memories. This is in line with theoretical frameworks of memory consolidation, which suggest that during encoding, the hippocampal index coordinates activity in cortical modules representing an experience.^15,19–21^ During consolidation (particularly during SWS), the brain rearranges its connectivity matrix to optimize information storage, and repeated reactivations of the hippocampal index promote intermodular connections to facilitate consolidation, thereby leading to greater network integration.^15^ Indeed, a recent study using rodent models provides complementary evidence for the importance of resting-state network integration for memory consolidation^82^ and work in humans shows that network integration supports the consolidation of motor sequence learning,^83^ as well as better working memory performance^27^ and other aspects of cognitive function.^84^

The findings from the current study point to the importance of NREM SWS for overnight emotional memory retention, particularly for the consolidation of negative stimuli. Much research suggests that emotionally laden experiences are better remembered than neutral ones following post learning nocturnal sleep^85–88^ or a daytime nap.^89^ Indeed, negative information appears to be preferentially retained following sleep, even at the expense of neutral elements.^34–36^ While much research traditionally implicates REM sleep in emotional memory processing,^62^ we did not observe any statistically significant relationship between overnight emotional memory retention and REM sleep expression in our sample. Other studies have suggested that SWS may also be involved in emotional memory consolidation. Research in rodent models suggests that reactivation of hippocampal-amygdala neuronal ensembles during NREM sleep supports contextual-threat learning.^90^ Greater SWS expression during a daytime nap^89^ or nocturnal sleep^91^ is linked to better retention of negative information. Studies using targeted memory reactivation (TMR), a technique that pairs prior learning with a sensory cue that is replayed during subsequent sleep, have shown that sensory cue replay during early NREM sleep (first 3h of sleep) benefits memory performance for associated negative stimuli,^92^ and that cue replay during SWS is associated with better visuospatial recall of negative pictures,^93^ as well as shorter reaction times for negative images.^94^ Despite the fact that we did not find any associations with REM sleep, it is likely that both NREM and REM sleep are important for emotional memory consolidation, with each potentially playing a distinct, but complementary role in sleep-dependent memory processing.^93,95^ Additionally, future work should investigate electrophysiological features of REM (e.g., theta power) which may be more sensitive than REM percentage of sleep time.

Results from the present data indicate that the positioning of the hippocampal formation in the resting-state functional network hierarchy contributes to overnight memory retention as well as the expression of key sleep oscillations critical for memory consolidation. While our data are derived from resting-state fMRI, and therefore are not a direct measurement of neuronal activity during NREM SWS, the findings are consistent with the literature and theories that implicate the importance of hippocampal involvement in directing network dynamics for memory consolidation. Evidence from animal models suggests that reactivation of hippocampal neuronal firing patterns during NREM SWS^18,96^ facilitates memory consolidation. Hippocampal sharp-wave ripples (SWRs) coordinate reactivation of neuronal ensembles involved in learning during wake^97,98^ and coincide with the simultaneous synchronization of cortical activity,^99^ potentially reflecting reactivation a ‘hippocampal index code’. In support of this theoretical framework, early studies in rodent models demonstrated that hippocampal CA1 place cells that extensively fired during waking behavior also extensively fired during subsequent sleep,^100^ and that the correlation pattern of place cell activity during NREM SWS mirrored the pattern observed while a behaving animal explored an environment.^18^ Since then, many studies have provided further evidence for hippocampal replay during sleep in animal models,^17,18,101,102^ with some more recent works even demonstrating the importance of selective reactivation for the consolidation of emotional memories.^103^ In human participants, data from fMRI has provided indirect evidence for memory-relevant hippocampal activation,^104,105^ and intracranial electrophysiological recordings in human patients have further shown that the precise temporal coupling between hippocampal SWRs, neocortical SOs, and thalamo-cortical sleep spindles reflects hippocampal-neocortical dialogue important for memory consolidation.^78,79^ In line with this literature, we found that greater hippocampal influence over functional network architecture was tied to stronger frontal SO-SP coupling. Altogether, research from multiple levels of analysis supports the role of the hippocampus in directing network activity to facilitate memory consolidation.

Greater hippocampal and amygdala influence over resting-state functional network organization was associated with better emotional memory retention. Importantly, these findings were specific to hippocampus and amygdala nodes, as occipital cortex, medial frontal gyrus, and parahippocampal nodes were not associated with emotional memory retention. CA determined that both the hippocampus and amygdala contribute unique as well as shared information to the overnight consolidation of emotional memories. While some work implicates sleep in gist abstraction and generalization,^106,107^ the present data may indicate that in individuals with more intrinsically influential MTL nodes, sleep aids in preserving pattern separation processes by stabilizing distinct internal representations of emotional events. Earlier work from our laboratory has shown that amygdala-hippocampal DG/CA3 coactivation during task performance supports successful emotional mnemonic discrimination.^9,108^ Recent work in mouse models has shown that optogenetically inhibiting neuronal reactivation during sleep leads to fear memory generalization due to an inability to discriminate between contexts.^103^ This may suggest that under normal conditions of neuronal reactivation, the integrity of an emotional event is preserved. Similarly, the present data indicate that participants with higher hippocampal or amygdala EC were able to better discriminate emotional lures from targets following overnight sleep. Altogether, these findings suggest greater hippocampal and amygdala influence over resting-state functional network architecture supports emotional memory fidelity over nocturnal sleep.

Greater hippocampal control over resting-state network information flow (BC) was specifically associated with emotional memory performance at immediate test, whereas amygdala mediation of information transmission was tied to overnight retention and delayed retrieval of emotional items. The dissociation of these MTL regions for different stages of memory processing was further supported by RF and ROC analyses, which implicated hippocampal centrality measures in memory acquisition (although with low predictive power) and different combinations of amygdala and hippocampal graph metrics as the most important features for overnight consolidation and delayed retrieval. These findings are in line with prior literature and known interactions between hippocampus and amygdala MTL subsystems that support emotional memory processing. While the hippocampal formation is implicated in memory encoding and processing, the amygdala is primarily associated with a modulation of consolidation. Through both direct and indirect anatomical connections,^32^ the amygdala modulates hippocampal representations by providing salience cues via stress hormone action and neuromodulatory influences that promote memory consolidation.^12,32^ Prior work from our research group has shown that basolateral amygdala (BLA) BOLD signal activity increases in response to emotional stimuli, whereas hippocampal DG/CA3 activity is enhanced only in response to correctly differentiating highly similar emotional events.^9,33^ Other studies have found similar results,^109,110^ demonstrating that hippocampal-amygdala activity supports successful retrieval of emotional memory,^11,111^ and that bidirectional communication between hippocampus and amygdala facilitates accurate emotional mnemonic discrimination.^108^ Therefore, the findings that hippocampus and amygdala graph metrics that assess the degree to which these nodes direct network activity are linked to distinct phases of memory processing are perhaps unsurprising. It will be interesting for future studies to ascertain how hippocampal-amygdala graph theoretical metrics dynamically evolve during memory encoding, consolidation, and retrieval.

Notably, findings of the present study were specific to stimuli of negative valence. Stimuli including negative content, in particular, have been shown to exhibit a sleep benefit.^34–36^ In fact, prior work has shown that the specific details of negative images are better remembered than those of their neutral or positive counterparts.^112,113^ This may be due to the emotional salience of negative elements leading to enhanced arousal during encoding which tags emotional events for preferential consolidation during sleep^114,115^ or to increased connectivity between amygdala, cortical areas, and other MTL structures.^112,116^ Although speculative, this may be beneficial from an evolutionary perspective. By prioritizing goal-relevant features of an experience, such as those eliciting a highly salient, emotional response or consequence, consolidation may have adaptive value by engraining dangerous or threating situations into memory for future avoidance, thereby yielding a greater evolutionary benefit.^117^ Taken together, this may partially explain why we only observed statistically significant relationships using LDI calculations for negative items.

The present analyses were conducted using a sample of cognitively unimpaired older adults. While age was not a significant predictor in any of our models, it is important to note that our age range was restricted to 60-85 years, thereby minimizing our ability to detect significant age effects. While we did not observe any effects of age in our sample, prior literature has demonstrated that these network measures change in both healthy and pathological aging. As compared to their younger counterparts, older adults exhibit a local breakdown of resting-state modular organization, with larger modules segregating into smaller modules that change topological composition.^25^ Increasing age is associated with reduced modularity of the control, attentional, limbic, and visual resting-state networks,^118,119^ as well as decreases in resting-state network segregation which predicts lower episodic memory factor scores.^120^ EC appears to be relatively stable across healthy aging^121^ but some work indicates that pathological aging, such as Alzheimer’s disease (AD), impacts EC mapping.^122–125^ Indeed, BC in some subcortical and cortical areas also change as a function of age and across the AD continuum.^126^ Age- and disease-associated changes in each of these graph theoretical metrics may have functional significance in the context of sleep-dependent memory processing, as the mechanisms underlying memory consolidation during sleep also change in healthy and abnormal aging and are linked to impaired overnight memory retention.^127,128^ Future work should examine these graph measures across wider age ranges, including in children and young adults, to determine the stability of these mechanisms and relevance for sleep-dependent memory consolidation, as well as to determine whether disruption of these metrics contributes to AD-related impairments in sleep-dependent memory processing.

While the present analyses revealed no statistically significant associations between SDB severity and measures of resting-state network architecture or emotional memory retention, it is well established that SDB impacts both MTL structure and function,^129,130^ as well as memory processing.^131,132^ Obstructive sleep apnea (OSA), one of the most common types of SDB, is a sleep disorder hallmarked by complete (apnea) or partial (hypopnea) upper airway collapse. The frequent cessations in breathing lead to sleep fragmentation, intermittent hypoxia, hypercarbia, increased inflammatory profiles, and oxidative stress which act in concert to drive neural damage and cerebrovascular injury.^133^ The estimated prevalence of OSA is as high as 38% in the general population, and it increases substantially to nearly 90% in older ages.^38,39^ OSA impacts hippocampal^134^ and amygdala^135^ resting-state functional connectivity, though impacts on network neuroscientific measures are less studied. However, some studies show that sleep deprivation^136,137^ and another common sleep disorder, insomnia,^138^ are linked to lower network modularity. OSA is additionally linked to memory deficits, and a recent study found adults with OSA exhibit impaired specific memory recognition for negative stimuli.^139^ The sample used in the current analyses consisted of cognitively intact older adults who exhibited varying degrees of SDB symptoms but were not actively treated. It is possible that we did not detect any significant effects of SDB due to the limited parametric range of disease severity in our sample, as most of the participants exhibited either no or mild SDB symptoms (as evidenced by AHI ≤ 15). Future work including a larger sample enriched for SDB with a greater range of symptom severity could allow for the investigation of clinical diagnostic subgroup analyses which may yield greater insight into OSA effects on resting-state network architecture and related emotional memory function.

This study has some limitations worth noting. The rsfMRI scan occurred at a separate timepoint from the overnight sleep study. While this was adjusted for in follow-up analyses, the graph metrics are not directly tied to task performance. Therefore, these measures reflect an intrinsic measure of functional network architecture. Additionally, some of the scans were collected using a protocol with a limited FOV and short scan time to ensure high-resolution MTL images while limiting participant burden. This comes at the cost of including some cortical regions and the ability to investigate dynamic changes. Recent advances in fMRI methods enable fast scan times while maintaining high resolution quality; future studies should utilize these methods.^140^ The sample size was relatively small and consisted primarily of Caucasian community-dwelling older adults from Orange County, CA, which may limit the generalizability of results to other populations. Another limitation is that the participants had varying degrees of SDB. While AHI was adjusted for in all models, SDB impacts on sleep expression,^60^ cognitive health,^141^ and neurobiology are complex.^142^ Additionally, our sample size for probing sleep oscillations in relation to graph theoretical metrics and memory was small. However, despite the limited sample size we were still able to observe meaningful relationships. Future research should include a larger, more diverse sample free of OSA symptoms to investigate these mechanisms. Given the lack of a wake-control group, we cannot ascertain whether these findings are truly sleep-dependent, however our associations with NREM SWS suggest that sleep does indeed play an important role in these relationships. There is the potential for a ‘first night effect’ on sleep architecture measures, and therefore the proportion of N3 used in mediation analyses may be a slight underestimate. However, the examination of individual differences rather than within-subject change is a strength in this regard. While the current study is correlative, this work has successfully identified potential mechanistic targets for intervention or manipulation, thereby laying a crucial foundation for future research to probe causal relationships between functional network architecture, sleep expression, and sleep-related memory processing. Altogether, despite these limitations, this unique approach provides novel insight into intrinsic brain network properties important for emotional memory processing and retention over a sleep period.

To the authors knowledge, this is the first study to apply graph theoretical techniques to probe central tenets of memory consolidation theory and the sleep oscillatory activity implicated in them. The use of the emotional mnemonic discrimination task afforded us the ability to probe valence-specific differences in the topological roles of key nodes of the MTL memory system. The findings from the current study provide an innovative analytical and conceptual framework for future investigation in both healthy and diseased populations.

## ACKNOWLEDGEMENTS

We would like to express our gratitude to all the participants who generously devoted their time to contribute to this study, without whom this research would not be possible. We would also like to acknowledge the laboratory technicians, specialists, and undergraduate research assistants who aided in participant recruitment and data collection. This research was supported by NIA grants F31AG074703 (to MGCF), F32AG074621 (to JNA), R21AG07955 (to BAM and MAY), R01AG053555 (to MAY), K01AG058353 (to BAM), and the American Academy of Sleep Medicine Strategic Research Award (to RMB and BAM).

## AUTHOR CONTRIBUTIONS

Conceptualization, M.G.C-F, M.A.Y, B.A.M; Methodology, M.G.C-F, J.N.A, R.F.B. D.E.B., H.N., N.S.S.; Software, J.C.J, A.D., D.E.B., H.N., N.S.S.,; Formal Analysis, M.G.C-F., J.N.A., S.K.; Investigation, M.G.C-F, D.E.B., A.D., I.Y.C., K.K.L., N.S.S.; Resources, A.B.N., R.M.B., M.A.Y., B.A.M.; Writing—Original Draft, M.G.C-F.; Writing—Review & Editing, M.G.C-F., J.N.A., R.F.B., S.K., D.E.B., I.Y.C., N.S.S., A.B.N., R.M.B., M.A.Y., B.A.M.; Visualization, M.G.C-F., J.N.A., S.K.; Supervision—R.M.B., M.A.Y., B.A.M, Project Administration—R.M.B., M.A.Y., B.A.M.; Funding Acquisition—M.G.C-F., R.M.B., M.A.Y., B.A.M.

