## Supplemental Tables and Figures for "Medial temporal lobe functional network architecture supports sleep-related emotional memory processing in older adults"

| Region | Total ROIs | Included ROIs | % Included |
|---|---|---|---|
| Frontal Lobe | 68 | 30 | 44.1% |
| Superior Frontal Gyrus | 14 | 2 | 14.3% |
| Middle Frontal Gyrus | 14 | 5 | 35.7% |
| Inferior Frontal Gyrus | 12 | 10 | 83.3% |
| Orbital Gyrus | 12 | 11 | 91.7% |
| Precentral Gyrus | 12 | 2 | 16.7% |
| Paracentral Lobule | 4 | 0 | 0% |
| Temporal Lobe | 56 | 50 | 89.3% |
| Superior Temporal Gyrus | 12 | 11 | 91.7% |
| Middle Temporal Gyrus | 8 | 8 | 100% |
| Inferior Temporal Gyrus | 14 | 10 | 71.4% |
| Fusiform Gyrus | 6 | 6 | 100% |
| Parahippocampal Gyrus | 12 | 11 | 91.7% |
| Posterior Superior Temporal Sulcus | 4 | 4 | 100% |
| Parietal Lobe | 38 | 2 | 5.3% |
| Superior Parietal Lobe | 10 | 0 | 0% |
| Inferior Parietal Lobe | 12 | 0 | 0% |
| Precuneus | 8 | 0 | 0% |
| Postcentral Gyrus | 8 | 2 | 25% |
| Cingulate Gyrus | 26 | 17 | 65.4% |
| Insular Gyrus | 12 | 12 | 100% |
| Cingulate Gyrus | 14 | 5 | 35.7% |
| Occipital Lobe | 22 | 16 | 72.7% |
| Medioventral Occipital Cortex | 10 | 10 | 100% |
| Lateral Occipital Cortex | 12 | 6 | 50% |
| Subcortical Regions | 36 | 36 | 100% |
| Amygdala | 4 | 4 | 100% |
| Hippocampus | 4 | 4 | 100% |
| Basal Ganglia | 12 | 12 | 100% |
| Thalamus | 16 | 16 | 100% |
| Total | 246 | 151 | 61.4% |

**Supplementary Table 1.** Areas covered by the Regions of Interest (ROI) from the Brainnetome Atlas included in the analysis mask for graph theoretical analyses.

| Brainnetome ROI Number | Region Label | Original Volume | Masked Volume | % in Mask |
|---|---|---|---|---|
| 13 | A10m_L | 1075 | 619 | 57.6 |
| 14 | A10m_R | 924 | 663 | 71.8 |
| 19 | A46_L | 805 | 549 | 68.2 |
| 20 | A46_R | 1019 | 567 | 55.6 |
| 22 | A9/46v_R | 946 | 689 | 72.8 |
| 27 | A10l_L | 794 | 466 | 58.7 |
| 28 | A10l_R | 1031 | 788 | 76.4 |
| 31 | IFS_L | 384 | 227 | 59.1 |
| 32 | IFS_R | 320 | 295 | 92.2 |
| 33 | A45c_L | 318 | 197 | 61.9 |
| 34 | A45c_R | 374 | 297 | 79.4 |
| 35 | A45r_L | 381 | 216 | 56.7 |
| 36 | A45r_R | 412 | 298 | 72.3 |
| 37 | A44op_L | 471 | 471 | 100.0 |
| 38 | A44op_R | 557 | 557 | 100.0 |
| 39 | A44v_L | 280 | 248 | 88.6 |
| 40 | A44v_R | 273 | 242 | 88.6 |
| 41 | A14m_L | 518 | 486 | 93.8 |
| 42 | A14m_R | 630 | 626 | 99.4 |
| 43 | A12/47o_L | 536 | 412 | 76.9 |
| 44 | A12/47o_R | 491 | 371 | 75.6 |
| 45 | A11l_L | 926 | 591 | 63.8 |
| 46 | A11l_R | 1210 | 880 | 72.7 |
| 47 | A11m_L | 637 | 462 | 72.5 |
| 48 | A11m_R | 732 | 492 | 67.2 |
| 50 | A13_R | 800 | 596 | 74.5 |
| 51 | A12/47l_L | 568 | 410 | 72.2 |
| 52 | A12/47l_R | 494 | 425 | 86.0 |
| 61 | A4tl_L | 434 | 374 | 86.2 |
| 62 | A4tl_R | 424 | 308 | 72.6 |
| 69 | A38m_L | 727 | 564 | 77.6 |
| 70 | A38m_R | 661 | 360 | 54.5 |
| 71 | A41/42_L | 505 | 468 | 92.7 |
| 72 | A41/42_R | 371 | 367 | 98.9 |
| 73 | TE1.0/TE1.2_L | 801 | 781 | 97.5 |
| 74 | TE1.0/TE1.2_R | 729 | 686 | 94.1 |
| 75 | A22c_L | 585 | 348 | 59.5 |
| 77 | A38l_L | 497 | 431 | 86.7 |
| 78 | A38l_R | 649 | 545 | 84.0 |
| 79 | A22r_L | 644 | 552 | 85.7 |
| 80 | A22r_R | 367 | 311 | 84.7 |
| 81 | A21c_L | 568 | 356 | 62.7 |
| 82 | A21c_R | 707 | 414 | 58.6 |
| 83 | A21r_L | 784 | 445 | 56.8 |
| 84 | A21r_R | 954 | 701 | 73.5 |
| 85 | A37dl_L | 619 | 335 | 54.1 |

| | | | | |
|---|---|---|---|---|
| 86 | A37dl_R | 726 | 503 | 69.3 |
| 87 | aSTS_L | 830 | 711 | 85.7 |
| 88 | aSTS_R | 1179 | 967 | 82.0 |
| 89 | A20iv_L | 317 | 232 | 73.2 |
| 90 | A20iv_R | 184 | 113 | 61.4 |
| 91 | A37elv_L | 336 | 268 | 79.8 |
| 92 | A37elv_R | 223 | 133 | 59.6 |
| 93 | A20r_L | 484 | 247 | 51.0 |
| 95 | A20il_L | 464 | 391 | 84.3 |
| 96 | A20il_R | 454 | 276 | 60.8 |
| 97 | A37vl_L | 407 | 294 | 72.2 |
| 98 | A37vl_R | 348 | 254 | 73.0 |
| 99 | A20cl_L | 499 | 423 | 84.8 |
| 103 | A20rv_L | 1017 | 693 | 68.1 |
| 104 | A20rv_R | 1111 | 559 | 50.3 |
| 105 | A37mv_L | 899 | 878 | 97.7 |
| 106 | A37mv_R | 801 | 788 | 98.4 |
| 107 | A37lv_L | 952 | 881 | 92.5 |
| 108 | A37lv_R | 890 | 758 | 85.2 |
| 109 | A35/36r_L | 175 | 163 | 93.1 |
| 110 | A35/36r_R | 143 | 124 | 86.7 |
| 111 | A35/36c_L | 153 | 112 | 73.2 |
| 112 | A35/36c_R | 159 | 113 | 71.1 |
| 113 | TL_L | 172 | 172 | 100.0 |
| 114 | TL_R | 134 | 134 | 100.0 |
| 115 | A28/34_L | 179 | 164 | 91.6 |
| 116 | A28/34_R | 120 | 60 | 50.0 |
| 117 | TI_L | 93 | 88 | 94.6 |
| 119 | TH_L | 148 | 146 | 98.6 |
| 120 | TH_R | 157 | 157 | 100.0 |
| 121 | rpSTS_L | 319 | 319 | 100.0 |
| 122 | rpSTS_R | 351 | 351 | 100.0 |
| 123 | cpSTS_L | 353 | 353 | 100.0 |
| 124 | cpSTS_R | 290 | 261 | 90.0 |
| 157 | A1/2/3tonIa_L | 649 | 461 | 71.0 |
| 158 | A1/2/3tonIa_R | 612 | 428 | 69.9 |
| 163 | G_L | 328 | 328 | 100.0 |
| 164 | G_R | 276 | 272 | 98.6 |
| 165 | vIa_L | 239 | 238 | 99.6 |
| 166 | vIa_R | 210 | 208 | 99.0 |
| 167 | dIa_L | 254 | 254 | 100.0 |
| 168 | dIa_R | 223 | 223 | 100.0 |
| 169 | vId/vIg_L | 261 | 261 | 100.0 |
| 170 | vId/vIg_R | 287 | 286 | 99.7 |
| 171 | dIg_L | 288 | 288 | 100.0 |
| 172 | dIg_R | 273 | 268 | 98.2 |
| 173 | dId_L | 392 | 392 | 100.0 |

| | | | | |
|---|---|---|---|---|
| 174 | dId_R | 308 | 308 | 100.0 |
| 178 | A24rv_R | 338 | 218 | 64.5 |
| 181 | A23v_L | 355 | 326 | 91.8 |
| 182 | A23v_R | 310 | 252 | 81.3 |
| 187 | A32sg_L | 636 | 636 | 100.0 |
| 188 | A32sg_R | 408 | 408 | 100.0 |
| 189 | cLinG_L | 492 | 481 | 97.8 |
| 190 | cLinG_R | 603 | 575 | 95.4 |
| 191 | rCunG_L | 785 | 676 | 86.1 |
| 192 | rCunG_R | 837 | 708 | 84.6 |
| 193 | cCunG_L | 613 | 520 | 84.8 |
| 194 | cCunG_R | 624 | 493 | 79.0 |
| 195 | rLinG_L | 760 | 751 | 98.8 |
| 196 | rLinG_R | 861 | 861 | 100.0 |
| 197 | vmPOS_L | 985 | 810 | 82.2 |
| 198 | vmPOS_R | 1033 | 810 | 78.4 |
| 199 | mOccG_L | 837 | 603 | 72.0 |
| 200 | mOccG_R | 858 | 530 | 61.8 |
| 201 | V5/MT+_L | 751 | 560 | 74.6 |
| 202 | V5/MT+_R | 807 | 593 | 73.5 |
| 205 | iOccG_L | 1024 | 735 | 71.8 |
| 206 | iOccG_R | 945 | 725 | 76.7 |
| 211 | mAmyg_L | 170 | 145 | 85.3 |
| 212 | mAmyg_R | 217 | 180 | 82.9 |
| 213 | lAmyg_L | 86 | 84 | 97.7 |
| 214 | lAmyg_R | 134 | 134 | 100.0 |
| 215 | rHipp_L | 574 | 568 | 99.0 |
| 216 | rHipp_R | 486 | 475 | 97.7 |
| 217 | cHipp_L | 588 | 585 | 99.5 |
| 218 | cHipp_R | 613 | 608 | 99.2 |
| 219 | vCa_L | 463 | 440 | 95.0 |
| 220 | vCa_R | 334 | 318 | 95.2 |
| 221 | GP_L | 324 | 287 | 88.6 |
| 222 | GP_R | 325 | 295 | 90.8 |
| 223 | NAC_L | 321 | 289 | 90.0 |
| 224 | NAC_R | 395 | 385 | 97.5 |
| 225 | vmPu_L | 322 | 295 | 91.6 |
| 226 | vmPu_R | 246 | 231 | 93.9 |
| 227 | dCa_L | 507 | 442 | 87.2 |
| 228 | dCa_R | 674 | 629 | 93.3 |
| 229 | dlPu_L | 408 | 408 | 100.0 |
| 230 | dlPu_R | 401 | 398 | 99.3 |
| 231 | mPFtha_L | 209 | 208 | 99.5 |
| 232 | mPFtha_R | 182 | 182 | 100.0 |
| 233 | mPMtha_L | 126 | 126 | 100.0 |
| 234 | mPMtha_R | 196 | 196 | 100.0 |
| 235 | Stha_L | 143 | 143 | 100.0 |

| | | | | |
|---|---|---|---|---|
| 236 | Stha_R | 150 | 150 | 100.0 |
| 237 | rTtha_L | 204 | 196 | 96.1 |
| 238 | rTtha_R | 217 | 180 | 82.9 |
| 239 | PPtha_L | 246 | 244 | 99.2 |
| 240 | PPtha_R | 217 | 216 | 99.5 |
| 241 | Otha_L | 237 | 232 | 97.9 |
| 242 | Otha_R | 166 | 163 | 98.2 |
| 243 | cTtha_L | 204 | 197 | 96.6 |
| 244 | cTtha_R | 147 | 146 | 99.3 |
| 245 | lPFtha_L | 358 | 358 | 100.0 |
| 246 | lPFtha_R | 273 | 273 | 100.0 |

**Supplementary Table 2.** A complete list of all ROIs included in the analysis mask with corresponding Brainnetome Region Labels, original ROI volumes, masked volumes, and the percentage of each ROI included in the mask.

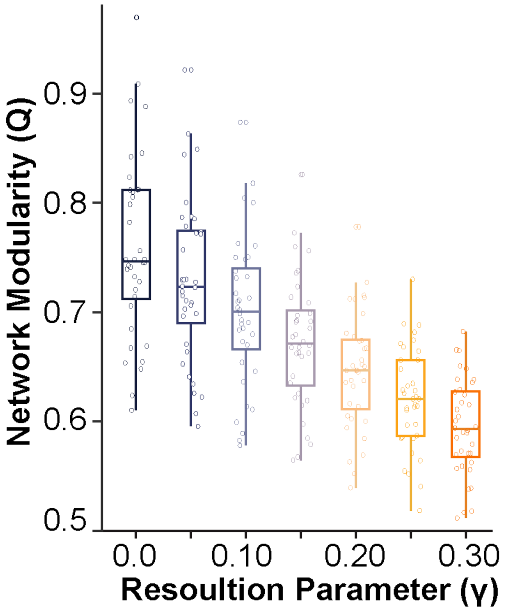


**Supplemental Figure 1.** Modularity across resolution parameters

| Network Measure | Kendall's $\tau_b$ (*p-value*) |
|---|---|
| Modularity ($\gamma$=0.10) | 0.199 *(0.089)* |
| Hippocampal Eigenvector Centrality | 0.037 *(0.754)* |
| Amygdala Eigenvector Centrality | 0.046 *(0.693)* |
| Hippocampal Betweenness Centrality | 0.032 *(0.785)* |
| Amygdala Betweenness Centrality | 0.016 *(0.892)* |

**Supplementary Table 3.** Associations between each network measure and mean motion for all valid resting-state fMRI scans.

| Network Measure | AHI<br>Pearson's r *(p-val)* | Time Below 90% Oxygen Saturation<br>Kendall's $\tau_b$ (*p-val*) |
|---|---|---|
| Modularity ($\gamma$=0.10) | 0.156 *(0.182)* | 0.185 *(0.121)* |
| Hippocampal Eigenvector Centrality | -0.206 *(0.077)* | -0.031 *(0.794)* |
| Amygdala Eigenvector Centrality | 0.222 *(0.057)* | 0.139 *(0.244)* |
| Hippocampal Betweenness Centrality | 0.027 *(0.817)* | 0.193 *(0.106)* |
| Amygdala Betweenness Centrality | 0.084 *(0.470)* | 0.036 *(0.763)* |

**Supplementary Table 4.** Associations between each network measure and indices of sleep disordered breathing severity. AHI represented log transformed AHI. AHI—Apnea-Hypopnea Index

| **Resolution Parameter (γ)** | **Pearson's *r* (p-value)** |
|---|---|
| 0 | 0.417 (0.012) |
| 0.05 | 0.411 (0.013) |
| 0.10 | 0.400 (0.016) |
| 0.15 | 0.377 (0.024) |
| 0.20 | 0.358 (0.032) |
| 0.25 | 0.346 (0.039) |
| 0.30 | 0.317 (0.060) |

**Supplementary Table 5.** Network modularity is positively associated with SWS expression across resolution parameters.

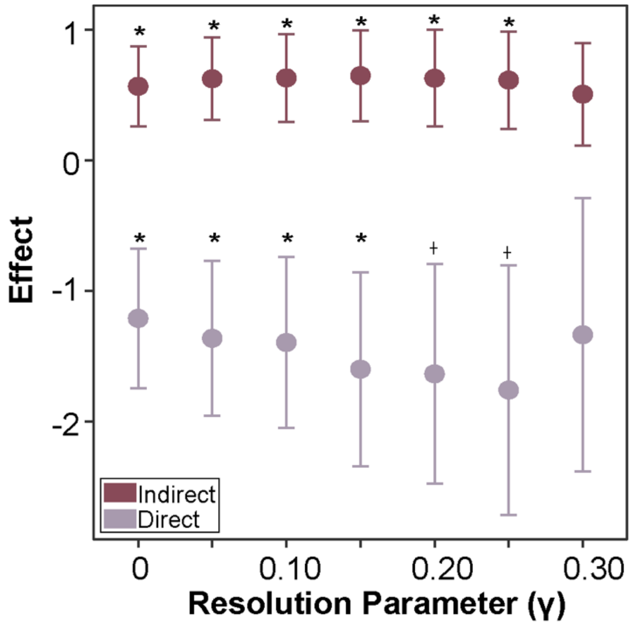


**Supplementary Figure 3.** Direct and indirect effects of network modularity on overnight negative memory retention across resolution parameters. *p≤0.05, †p<0.10

| | Immediate Test | | | Delayed Test | | |
|---|---|---|---|---|---|---|
| | Negative | Neutral | Positive | Negative | Neutral | Positive |
| LSim LDI | 0.69 ± 0.17 | 0.72 ± 0.20 | 0.55 ± 0.21 | 0.47 ± 0.22 | 0.44 ± 0.17 | 0.45 ± 0.18 |
| Target HR | 0.90 ± 0.07 | 0.91 ± 0.07 | 0.89 ± 0.09 | 0.85 ± 0.12 | 0.80 ± 0.14 | 0.80 ± 0.14 |
| LSim Lure CR | 0.78 ± 0.15 | 0.79 ± 0.15 | 0.63 ± 0.21 | 0.60 ± 0.22 | 0.64 ± 0.15 | 0.64 ± 0.18 |
| LSim Lure FA | 0.23 ± 0.14 | 0.22 ± 0.14 | 0.37 ± 0.20 | 0.40 ± 0.22 | 0.36 ± 0.15 | 0.36 ± 0.17 |

Abbreviations: LSim—Low Similarity; LDI—Lure Discrimination Index; HR—Hit Rate; CR—Correct Rejection Rate; FA—False Alarm Rate

**Supplementary Table 6.** Behavioral data for low similarity lure condition from the E-MDT at immediate and delayed testing timepoints.

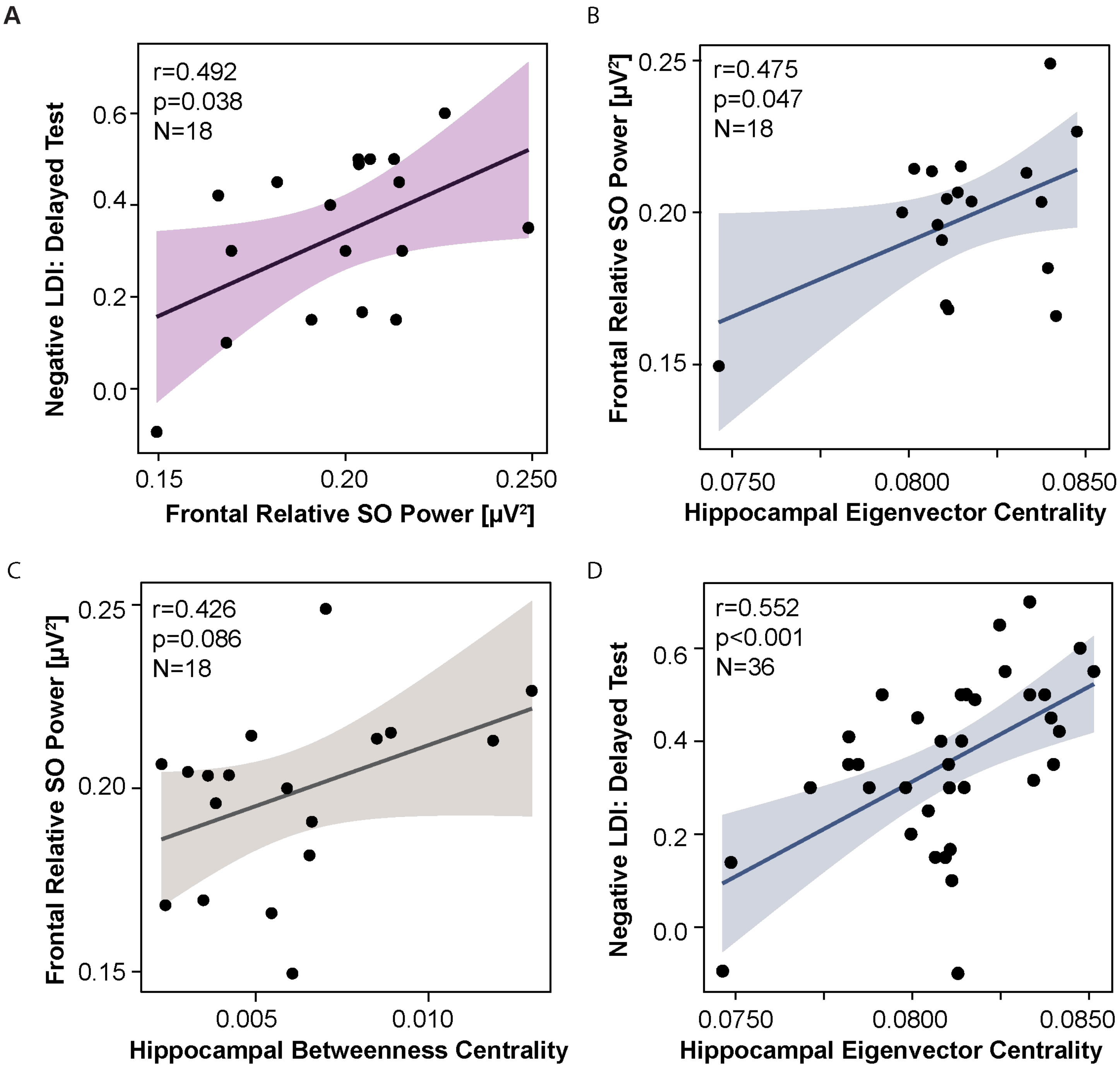


**Supplemental Figure 4.** Frontal slow oscillation power is tied to next-day emotional memory retrieval and measures of hippocampal centrality. (A) Greater frontal relative SO power is associated with better next-day retrieval of emotional memories. (B) Greater hippocampal eigenvector centrality is associated with greater frontal relative slow oscillation power. (C) Greater hippocampal betweenness centrality is associated with greater frontal relative SO power. (D) Greater hippocampal eigenvector centrality is associated with better next-day retrieval of emotional memories. Abbreviations: r—Pearson's correlation; p—p-value; N—sample size; LDI—Lure Discrimination Index; SO—slow oscillation

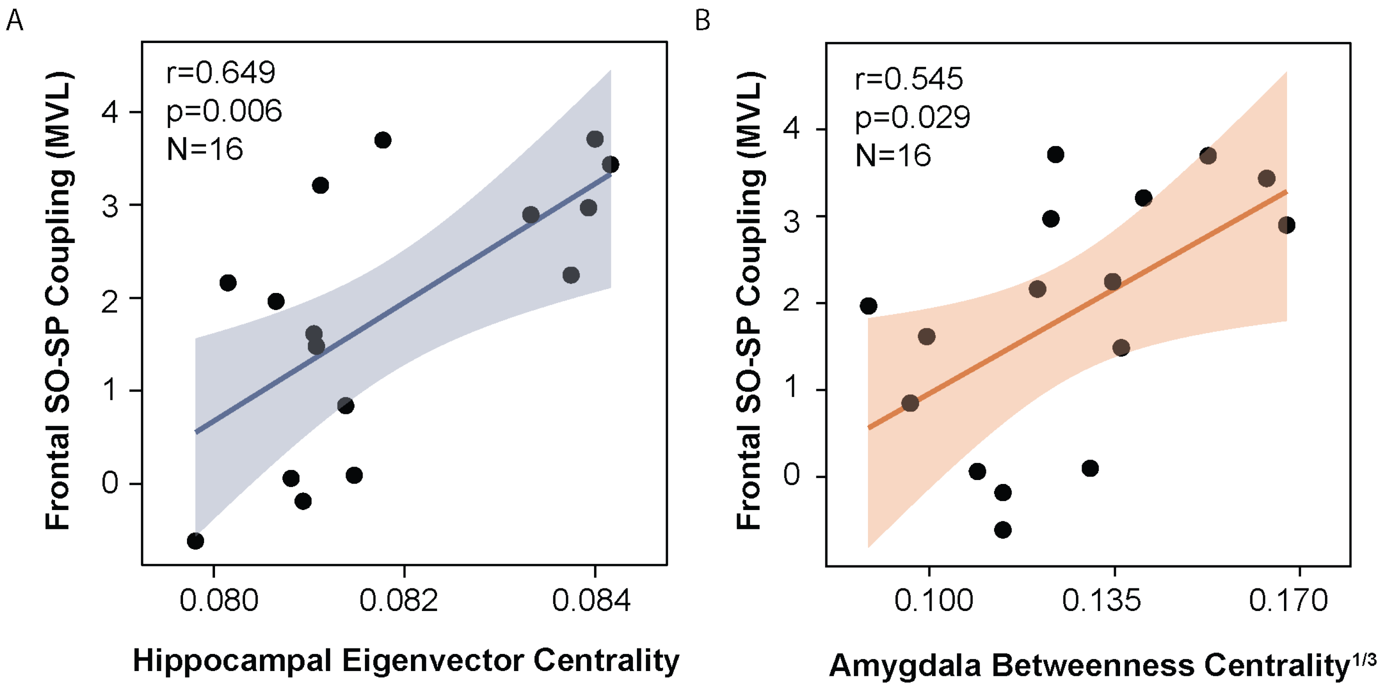


**Supplemental Figure 5.** Slow oscillation fast sleep spindle coupling are associated with measures of medial temporal lobe centrality. (A) Greater hippocampal eigenvector centrality is associated with greater phase amplitude coupling between slow oscillations and fast sleep spindles. (B) Greater amygdala betweenness centrality is associated with greater phase amplitude coupling between slow oscillations and fast sleep spindles. Abbreviations: r—Pearson's correlation; p—p-value; N—sample size; SO—slow oscillation; SP—fast sleep spindle; MVL—Mean Vector Length. Note: Amygdala betweenness centrality is cube-root transformed.
